# Mitochondrial complementation occurs in rat hippocampal axons and supports the synaptic vesicle cycle

**DOI:** 10.1101/2022.07.06.499030

**Authors:** Jason D. Vevea, Edwin R. Chapman

## Abstract

Mitochondria exert powerful control over cellular physiology, contributing to ion homeostasis, energy production, and metabolite biosynthesis. Mitochondrial trafficking and function are vital to neurons, with organelle impairment or altered morphology observed in every neurodegenerative disorder studied. While this organelle and its biosynthetic products are critical for cellular function, decreased output and/or byproducts (e.g., free radicals), can be harmful, and organelle quality control (QC) mechanisms are required to maintain function and prevent a cascade of damage. Owing to its length and general lack of biosynthetic machinery, the axon is particularly sensitive to damage and there is little consensus regarding the details of mitochondrial QC mechanisms in this compartment. Here we investigate the basal, unstressed behavior of axonal mitochondria, focusing on mitochondrial trafficking and fusion to better understand potential QC mechanisms. We observed size and redox asymmetry of mitochondrial traffic in the axon, suggestive of an active QC mechanism. Importantly, we demonstrate, in detail, biochemical complementation of axonal mitochondria. Upon disruption of mitochondrial fusion, we observed an altered synaptic proteome, presynaptic calcium dyshomeostasis, decreased levels of exocytosis, and a reduction in synaptic vesicle recruitment from the reserve pool during extended stimulation. These results support an active mitochondrial trafficking and fusion related QC process that supports presynaptic physiology.

- **Anterograde trafficked mitochondria are larger and relatively more reduced than retrograde trafficked mitochondria**
- **Anterograde mitochondria fuse with and complement resident, stationary, axonal mitochondria**
- **Loss of mitofusin 2 (MFN2) mediated mitochondrial fusion leads to alterations in the synaptic vesicle cycle and decreased reserve pool mobilization**

## Introduction

Mitochondria aid numerous cellular functions by supporting energy production (adenosine triphosphate, ATP), metabolite biosynthesis (lipids, amino acids, nucleotides, heme), ion homeostasis (K^+^, Ca^2+^), and the response to cellular stress **(Spinelli & Haigis, 2018)**. Mitochondria are morphologically dynamic, and differences in morphology between cell types may reflect specialized functions **(Eisner, Picard, & Hajnoczky, 2018)**. Mitochondrial morphology can even differ dramatically within a single cell. For example, neurons often consist of three compartments: a soma or cell body that houses the nucleus and most biosynthetic machinery, dendrites that receive chemical signals, and an axon that secretes chemical signals (exocytosis). Mitochondria form an interconnected web-like tubular network in the soma, a more parallel interconnected tubular network in dendrites, and shorter, seemingly disconnected mitochondrial units in axons **(Faitg et al., 2021; Lewis, Kwon, Lee, Shaw, & Polleux, 2018)**. Mitochondrial size and connectedness directly correlate with the Ca^2+^ buffering capacity of the mitochondrion, and these various morphologies throughout the neuron may reflect Ca^2+^ buffering differences **(Lewis et al., 2018)**. So, while mitochondria in the axon likely have a lower overall capacity to buffer Ca^2+^ relative to other neuronal compartments, the shorter and less contiguous axonal network of mitochondria may be particularly sensitive to perturbation. Indeed, presynaptic Ca^2+^ influences numerous steps during exocytosis and endocytosis of synaptic vesicles (SV) **(Chanaday, Cousin, Milosevic, Watanabe, & Morgan, 2019)**, and Ca^2+^ buffering by mitochondria regulates presynaptic Ca^2+^ and neurotransmitter release **(Billups & Forsythe, 2002)**.

Axonal mitochondria traffic long distances from the soma, their source of biogenesis, and must be maintained millimeters away from the nucleus **(Saxton & Hollenbeck, 2012)**. There is no consensus for how quality control (QC) over the mitochondrial network in axons is exerted. Mitochondrial QC is the term for cellular mechanisms related to the maintenance of mitochondrial function in the face of accumulated organelle damage over time. Axonal QC mechanisms targeting less functional or damaged mitochondria could happen locally in the axon or in the soma after a retrograde trafficking event. Mitochondria that are targeted for QC could be disposed of in the axon through autophagic mechanisms (mitophagy), restored through homotypic mitochondrial fusion with a higher functioning mitochondrion (complementation), or retrogradely transported back to the soma for mitophagy or complementation. Trafficking of mitochondria in axons is expected to be important for axonal function and neuronal health, because altered mitodynamics are found in many, if not all neurodegenerative diseases and disease models examined; importantly, this is also the case for sporadic cases **(Chen & Chan, 2009; Schon & Przedborski, 2011)**, but see also **(Area-Gomez, Guardia-Laguarta, Schon, & Przedborski, 2019)**.

The existence of axonal QC mechanisms regulating the asymmetric traffic of mitochondria remains the subject of debate. In unstressed axons, there is evidence both for **(Mandal et al., 2021; Miller & Sheetz, 2004)**, and against **(Suzuki, Hotta, & Oka, 2018; Verburg & Hollenbeck, 2008)** mitochondrial trafficking asymmetry, indicative of axonal trafficking QC. Indeed, there may not be a need for a trafficking QC mechanism in the axon if mitochondrial biogenesis **(Amiri & Hollenbeck, 2008; Kuzniewska et al., 2020)** and degradation (mitophagy) **(Ashrafi, Schlehe, LaVoie, & Schwarz, 2014)** occur locally. However, local biogenesis might be limited, and axonal mitophagy only occurs during extreme (likely non-physiological) stress. In contrast, low (nM) concentrations of mitochondrial toxins **(Cai, Zakaria, Simone, & Sheng, 2012; M. Y. Lin et al., 2017)**, and in specific disease models **(Ebrahimi-Fakhari et al., 2016; Zheng et al., 2019)**, increased mitochondrial retrograde traffic has been reported. Moreover, many key mitochondrial proteins are exceptionally long-lived **(Bomba-Warczak, Edassery, Hark, & Savas, 2021)**. In fact, mitophagy and autophagy might be completely dispensable for mitochondrial homeostasis in axons **(T. H. Lin et al., 2021)**. Interestingly, the phenomenon of mitochondrial trafficking asymmetry is evolutionarily conserved as yeast **(Higuchi et al., 2013; McFaline-Figueroa et al., 2011)** and stem cells **(Katajisto et al., 2015)** are able to segregate low from high functioning mitochondria via a QC mechanism that is critical for lifespan and maintaining stemness, respectively.

Here we examine axonal mitochondrial morphology, trafficking, and redox status, in unstressed neurons, using fluorescent probes (e.g., mitochondrial matrix targeted roGFP (mito-roGFP) **(Dooley et al., 2004; Hanson et al., 2004)**) and microfluidic devices. The relative redox status of the mitochondrial matrix is used as a proxy for fitness of the organelle, with more reduced ratio relating to better fitness **(McFaline-Figueroa et al., 2011)**. We observed size and redox mitochondrial trafficking asymmetry which are indicative of an axonal QC mechanism. We also observed frequent mitochondrial fusion and content mixing in axons, and using mito-roGFP, we provide evidence for redox complementation during axonal mitochondrial fusion. We then asked whether this mitochondrial fusion and complementation are critical for the function of the axon, i.e., the SV cycle. We addressed this question via knockdown (KD) of mitofusin 2 (MFN2), an outer mitochondrial membrane bound GTPase that mediates mitochondrial fusion in neurons **(Eura, Ishihara, Yokota, & Mihara, 2003)**. We then interrogate the SV cycle and presynaptic Ca^2+^ homeostasis using a SV targeted pHluorin (vGlut1-pHluorin) **(Voglmaier et al., 2006)** and the new far-red calcium sensor HTL-JF646-BAPTA-AM **(Deo, Sheu, Seo, Clapham, & Lavis, 2019)** targeted to SVs using synaptophysin-HaloTag (SYP-HT) **(Bradberry & Chapman, 2022)**. We find that when mitochondrial fusion is impaired, presynaptic function is disrupted. There are fewer SV proteins, altered endocytosis, decreased total exocytosis, a smaller recycling pool of SVs, and faster presynaptic Ca^2+^ transients.

## Results

### Asymmetric trafficking of mitochondria in axons

We used dissociated rat hippocampal neurons to study axonal mitochondrial morphology and trafficking parameters. Before focusing on axonal mitochondria, we first corroborated morphological differences between mitochondria in the soma (nuclear), dendrite (post-synaptic), and axonal (pre-synaptic) compartments (Figure 1a) **(Faitg et al., 2021; Lewis et al., 2018)**. Dendrites have spines and a large tapering shaft diameter, while axons are smooth, do not taper, and have a smaller diameter **(Craig & Banker, 1994)**. Clear dendritic (right neurite) and axonal (left neurite) morphological characteristics are revealed by cytosolic mCherry (Figure 1a, magenta box). Using mitochondrial targeted green fluorescent protein (mito-GFP) (Figure 1a, green box), we confirm that mitochondria in dendrites are long and overlapping, while axonal mitochondria are shorter and evenly spaced apart **(Faitg et al., 2021; Lewis et al., 2018)**. We then looked at trafficking using mitochondrial-targeted photo-activatable GFP (PA-GFP) **(Patterson & Lippincott-Schwartz, 2002)**. Photo-activation of the soma allowed us to monitor new mitochondrial trafficking events to neurites (Figure 1b). Mitochondria that entered the axon were small and unitary in size and showed processive movement throughout the image series (30 min). In contrast, mitochondria entered dendrites slowly and did not traffic far, appearing to immediately fuse with the dendritic mitochondrial network, similar to a previous report **(Overly, Rieff, & Hollenbeck, 1996)**. To further examine mitochondrial trafficking in the axon, we cultured primary neurons in microfluidic devices that separate the growth of axons from dendrites and the soma (Figure 1c; Video 1). Microfluidic isolation allowed axon vs somato-dendritic mitochondria counterstaining using different MitoTracker dyes. We then imaged axons in the middle of the 450-micron microfluidic channel to monitor retrograde (defined as MitoTracker Red +; MTR+) and anterograde (defined as MitoTracker Green +; MTG+) events. We found anterograde trafficked mitochondria were longer (Figure 1d) and more processive (Figure 1e) than retrograde trafficked mitochondria. The average speed of anterograde mitochondria was faster as well (Figure 1f). Together, these data establish the existence of mitochondrial trafficking asymmetry in axons, which may be related to an organelle QC mechanism.

**Figure 1.**
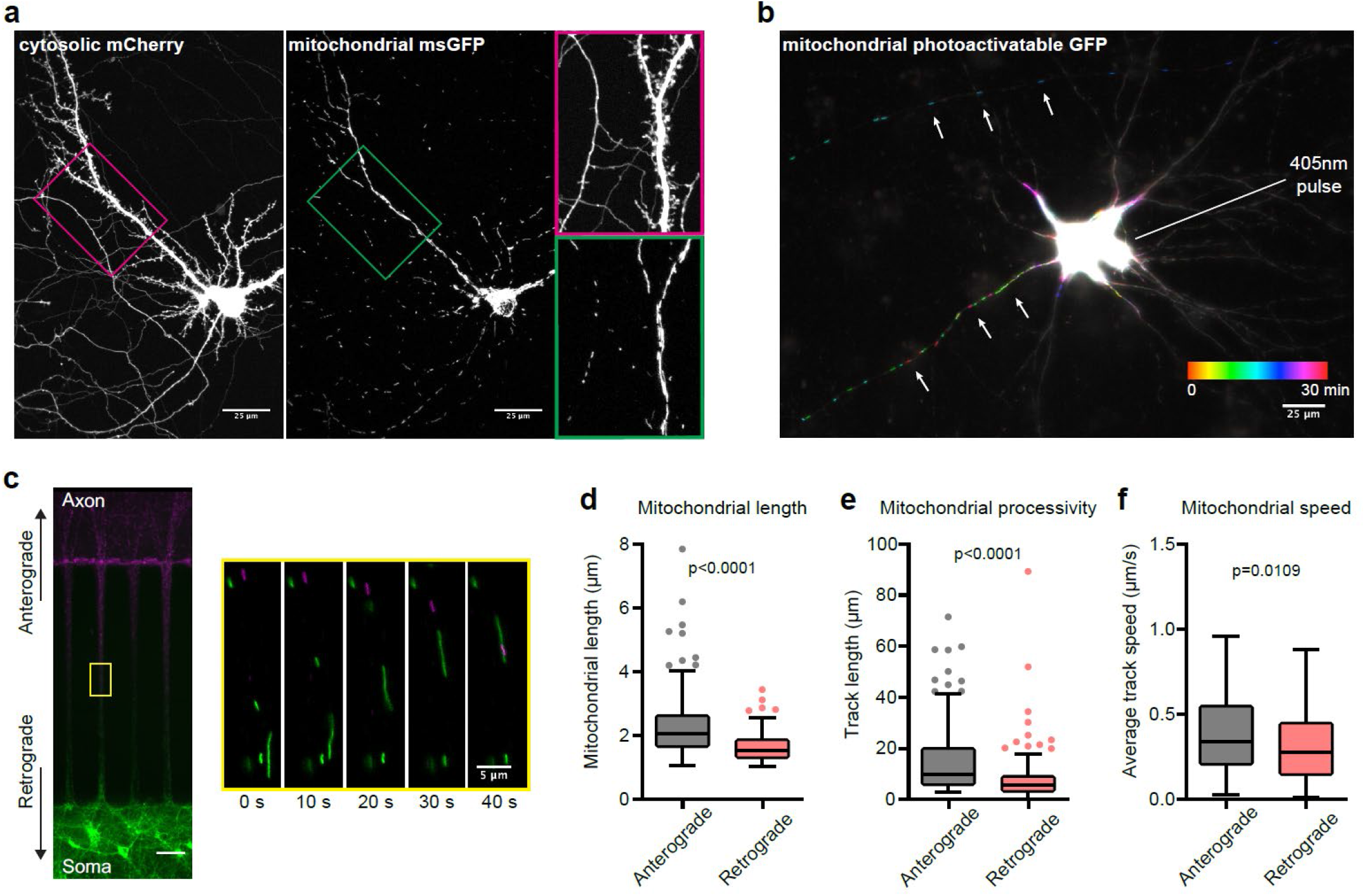
Asymmetric trafficking of mitochondria in axons. **a**) Maximum projection of a live-cell, confocal z-stack from a 16 DIV neuron co-transfected with cytosolic mCherry and mitochondrial matrix targeted monomeric-superfolder green fluorescent protein (msGFP). Scale bar: 25 µm. Magenta and green boxes outline zoomed-in areas on the right, highlighting the distinct morphologies of axonal and dendritic mitochondria. **b**) Temporal color-coded max projection of a 15 DIV neuron expressing mitochondrial matrix targeted photo-activatable GFP (PA-GFP). Mitochondrial soma egress was monitored after photoactivation by imaging every minute for 30 total minutes. Scale bar: 25 µm. **c**) Representative images of 14 DIV neurons, grown in SND450 (Xona) microfluidics, stained with MitoTracker Green FM (MTG) and MitoTracker Red CMXRos (MTR) on the soma and axon side, respectively. Scale bar: 50 µm. The yellow square marks the area enlarged on the right, showing delineated mitochondrial trafficking events at the indicated time points. Scale bar: 5 µm. Mitochondria imaged in this area have traveled at least 200 µm as microchannels in microfluidic are 450 µm long. **d**) Mitochondria were segregated into anterograde (black) or retrograde (red) groups based on MitoTracker dye labeling (green vs red) and direction of motility in the microchannel (see also Materials and Methods). Mitochondrial length quantified from middle of microchannel, anterograde (2.07 µm [95% Confidence Interval (CI) 1.88 - 2.15]) and retrograde (1.53 µm [95% CI 1.46 - 1.62]). Values are medians with 95% CI representing error, Mann-Whitney test p < 0.0001, detailed statistics in Figure 1 – source data 1. **e**) Track lengths (processive movement or un-paused motility), anterograde (black) (9.64 µm [95% CI 7.82 - 12.35]) and retrograde (red) (5.67 µm [95% CI 4.64 - 6.06]), segregated as in (d). Pause is defined as 3 consecutive images (15 sec) where organelle does not move. Values are medians with 95% CI representing error, Mann-Whitney test p < 0.0001, detailed statistics in Figure 1 – source data 2. **f**) Average track speed values (track distance over time) plotted by anterograde (black) or retrograde (red) direction, segregated as in (d). Anterograde (0.34 µm/s [95% CI 0.30 - 0.39]) and retrograde (0.28 µm/s [95% CI 0.24 - 0.31]) mitochondrial trafficking speeds are plotted. Values are medians with 95% CI representing error, Mann-Whitney test p = 0.0109, detailed statistics in Figure 1 – source data 3. Graphs in (d-f) are Tukey box and whisker plots with n > 160 for each group from at least three independent experiments.

### Anterogradely trafficked mitochondria are more reduced and complement resident axonal mitochondria

We next examined the redox status and fusion rates of axonal mitochondria using microfluidics, mitochondrial-targeted redox sensitive GFP (roGFP) **(Dooley et al., 2004; Hanson et al., 2004)**, and MitoTracker dyes. We transfected mito-roGFP into neurons, grew them in microfluidics, and imaged the axon chamber, just above the microchannel, for 5 min (Figure 2a, magenta box). Anterograde and retrograde trafficking mitochondria were identified, and their relative redox ratio quantified. We found that anterograde trafficking mitochondria had a more reduced mitochondrial matrix than stationary or retrograde trafficking mitochondria (Figure 2b), and non-normalized ratios are included in the Supplement (Figure 2 – Figure Supplement 1a). Because axon mitochondrial traffic asymmetry has been controversial, we again measured mitochondrial length, and again find anterograde mitochondria are longer (Figure 2c). The difference here is significant but slightly less than in Figure 1d, possibly because this experiment does not contain a counter stain to distinguish, without ambiguity, somato-dendritic derived anterograde mitochondria vs retrograde mitochondria. During our image series we identified several fusion, or complementation, events. In Figure 2d and Video 2 we show a representative example of an anterograde trafficking mitochondria fusing with a stationary axonal mitochondrion of a more oxidized status. The fused mitochondrion moved a small distance, the redox ratio equilibrated, and then the mitochondrion divided again. Values of redox ratio and length are included in the Table below the image frames. Axonal fusion events interested us greatly and so we monitored the rate of fusion of anterograde trafficked mitochondria by labeling neurons grown in a microfluidic device as in Figure 1, then imaging at hour timepoints after labeling (Figure 2e; Video 3). We monitored the rate of newly trafficked mitochondria into the axon chamber every hour, over four hours, by imaging at the cyan box shown in Figure 2e. Somato-dendritic mitochondria (MTG+) trafficked into the axon chamber at approximately 3.5% of total mitochondrial signal, per hour (Figure 2f). Approximately half (40% to 60%) of these newly trafficked mitochondria fuse with resident axonal mitochondria (Figure 2g). Lastly, we measured the rate of fusion among resident axonal mitochondria (MTR+) and found that they fuse (colocalize) with newly trafficked mitochondria (MTG+) at a rate of about 2% per hour (Figure 2h), this value represents the complementation rate. These results indicate that not only is there a physical asymmetry (Figure 1), there is also a functional (redox state) asymmetry regarding mitochondrial traffic in the axon; new, larger, and more reduced mitochondria complement axonal resident mitochondria.

**Figure 2.**
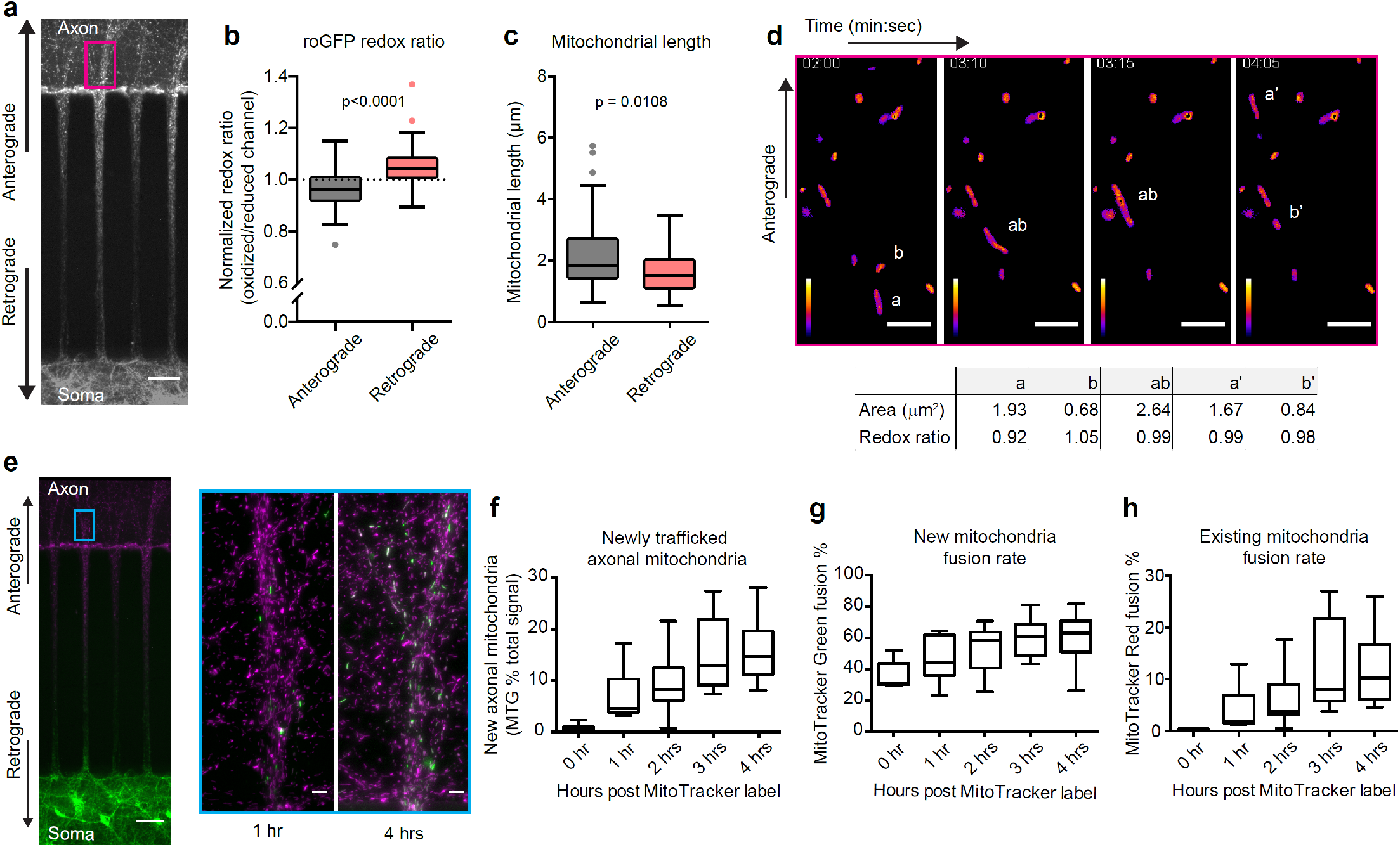
Anterograde mitochondria are more reduced and complement resident axonal mitochondria. **a**) Representative images of neurons grown in SND450 (Xona) microfluidics and transfected with mitochondrial matrix targeted redox-sensitive GFP (roGFP). Analysis in 2b-d was limited to fields of view (FOV) on the axon side near microchannels, denoted with the magenta rectangle. Scale bar 50 microns. **b**) Anterograde (black) or retrograde (red) trafficking mitochondria were grouped as in Figure 1 and redox ratio was calculated as described in Materials and Methods, then normalized to the average redox ratio of the population of mitochondria in the FOV. The redox ratio values were 0.96 [95% CI 0.94 - 0.97] (oxidized to reduced) for anterograde and 1.04 [95% CI 1.02 - 1.06] (oxidized to reduced) for retrograde. Values are medians with 95% CI representing error, Mann-Whitney test p < 0.0001, detailed statistics in Figure 2 – source data 1. **c**) As in Figure 1d, we compared the length of anterograde (black) trafficked mitochondria to retrograde (red) trafficked mitochondria. Anterograde trafficked mitochondria were again longer than retrograde, with values of (1.84 µm [95% CI 1.63 - 2.27]) and retrograde (1.52 µm [95% CI 1.19 - 1.95]). Values are medians with 95% CI representing error, Mann-Whitney test p = 0.0108, detailed statistics in Figure 2 – source data 2. Charts (2b and 2c) are Tukey box and whisker plots, with n > 50 for each group from at least three independent experiments. **d**) Representative images from an image series of axonal mito-roGFP. Low redox ratio mitochondria (*a*) can be seen moving in the anterograde direction toward stationary mitochondria (*b*). These mitochondria fuse and the redox ratio equilibrates as the now single mitochondrion (*ab*) continues in the anterograde direction. After a short distance it divides and a small piece, approximately the size of the original (*b*) mitochondrion, is left in a new position (*b’*). The rest of the mitochondrion (*a’*) continues in the anterograde direction. A table with size and redox ratio values is listed below the images. White scale bar: 5 µm, Fire LUT scale bar represents intensity: white = highest ratio, purple = lowest ratio. **e**) Representative image (left) of neurons grown (13 DIV to 16 DIV) in a microfluidic device as in 1c. Analysis of data in panels f-h was limited to fields of view (FOV) on the axon side near microchannels, denoted with the cyan rectangle. Scale bar was 50 µm (left) and 5 µm for enlarged areas. Representative images of 1 hour and 4 hours post MitoTracker labeling, displaying somato-dendritic mitochondrial traffic into axons and fusion with resident axonal mitochondria. **f**) A measure of newly trafficked axonal mitochondria, defined by MTG signal as a percentage of total fluorescent signal (MTG/(MTR + MTG)) at hour intervals post MitoTracker label. Newly trafficked mitochondria, account for 4.59% [95% CI 3.8% - 10.72%] of total mitochondria at 1 hour, 8.28% [95% CI 6.25% - 12.4%] at 2 hours, 12.9% [95% CI 9.13% - 21.91%] at 3 hours, and 14.71% [95% CI 11.19% - 19.32%] at 4 hours. Values are medians with 95% CI representing error, detailed statistics in Figure 2 – source data 3. **g**) A measure of the fraction of new mitochondria (visualized with MTG) that fuse with resident axonal mitochondria, defined as the percentage of MTG signal that colocalized with MTR signal. The fraction of newly trafficked mitochondria that undergo fusion is 43.9% [95% CI 35.8% - 61.9%] of new mitochondria at 1 hour, 58.2% [95% CI 42.9% - 63.4%] at 2 hours, 61.2% [95% CI 48.7% - 68.4%] at 3 hours, and 62.9% [95% CI 56.6% - 68.7%] at 4 hours. Values are medians with 95% CI representing error, detailed statistics are in Figure 2 – source data 4. **h**) Fraction of resident axonal mitochondria (visualized with MTR) that fuse with newly trafficked mitochondria, defined as the percentage of MTR that colocalized with MTG. These values are: 1.95% [95% CI 1.5% - 7.7%] of new mitochondria at 1 hour, 3.8% [95% CI 3.1% - 8.2%] at 2 hours, 8% [95% CI 5.8% – 21.7%] at 3 hours, and 10.2% [95% CI 6.4% - 15.7%] at 4 hours. Values are medians with 95% CI representing error, detailed statistics in Figure 2 – source data 5. Charts (2f, 2g, and 2h) are Tukey box and whisker plots with n = 5 for time 0 hour, and n > 15 for each other group, from at least three independent experiments.

**Figure 2 Figure supplement 1.**
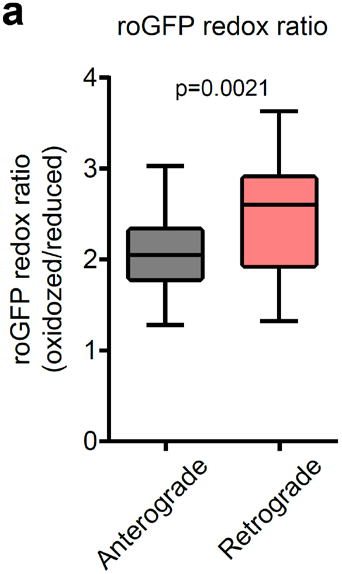
**a**) Non-normalized values for the data presented in Figure 2b. The redox ratio values are 2.05 [95% CI 1.95 - 2.25] (oxidized to reduced) for anterograde (black) and 2.6 [95% CI 2.18 - 2.83] (oxidized to reduced) for retrograde (red). Values are medians with 95% CI representing error, Mann-Whitney test p = 0.0021, detailed statistics in Figure 2 Figure supplement 1 – source data 1. Chart is a Tukey box and whisker plot.

### Mitochondrial fusion supports synaptic vesicle recycling

We next determined whether loss of mitochondrial fusion influences the synaptic vesicle cycle using a SV targeted pHluorin (vGlut1-pHluorin) **(Voglmaier et al., 2006)** as a read-out for exo- and endocytosis. Superecliptic pHluorin (referred to as pHluorin hereafter) is a pH sensitive GFP mutant with a pK of ∼7.1 **(Sankaranarayanan, De Angelis, Rothman, & Ryan, 2000)**. Upon SV exocytosis, pHluorin dequenches (i.e., the fluorescence increases), and during endocytosis and reacidification, requenches (i.e., the fluorescence decreases) (Figure 3d), allowing us to measure exocytosis and endocytosis. Mitochondrial fusion in vertebrates is mediated in part by the large outer membrane bound GTPases mitofusin 1 and 2 **(Chen et al., 2003)**. We disrupted mitochondrial fusion by knocking down (KD) the main mitofusin isoform (MFN2) in neurons **(Eura et al., 2003)**. With lentivirus shRNA targeted toward MFN2, we achieved stable and consistent MFN2 KD to approximately 20% that of control conditions (Figure 3a-b). In MFN2 KD neurons, mitochondria appeared swollen and less tubular (Figure 3c). We then investigated the SV cycle by monitoring the SV pHluorin response in CTRL and MFN KD neurons; an illustration of SV pHluorin response upon exocytosis is shown in Figure 3d. We measured the rate of endocytosis, as well as the readily releasable pool (RRP), the total reserve pool (RP), and total SV pool, using established protocols **(Li et al., 2005)**. Briefly, 40 action potentials (40 AP) drive the release of the RRP, while 900 AP drive the release of the RP. Endocytosis occurs during exocytosis, therefore, to accurately measure exocytosis, a V-ATPase inhibitor like folimycin is required to inhibit SV reacidification. Finally, fluorescence is normalized to bath application of NH4Cl, which is used to dequench all pHluorin at the synapse and measure total potential signal. The average traces for RRP and RP depleting stimuli are charted in Figure 3e. First, we found that the rate of endocytosis (tau; τ) was faster at MFN2 KD synapses than CTRL KD synapses (Figure 3f). The NH_4_Cl normalized RRP without folimycin (Figure 3 – Figure supplement 1a) and with (Figure 3g) was unchanged between CTRL KD and MFN2 KD synapses; however the size of the RP was significantly lower in MFN2 KD synapses (Figure 3h). The non-NH_4_Cl normalized RRP, with and without folimycin are included in Figure 3 – Figure supplement 1b-c, and show that absolute exocytosis is decreased by approximately half at the MFN2 KD synapse. The total vesicle pool, as measured by total vGlut1-pHluorin fluorescence during bath application of NH_4_Cl, was approximately half in MFN2 KD neurons relative to CTRL KD neurons (Figure 3i). We plotted the rate of endocytosis versus the total pHluorin measured SV pool but did not observe any obvious relationship (Figure 3 – Figure supplement 1d). These experiments show that mitochondrial fusion is necessary for the maintenance of normal rates of endocytosis, the absolute degree of exocytosis, mobilization of the RP during intense stimulation, and potentially for SV number.

**Figure 3.**
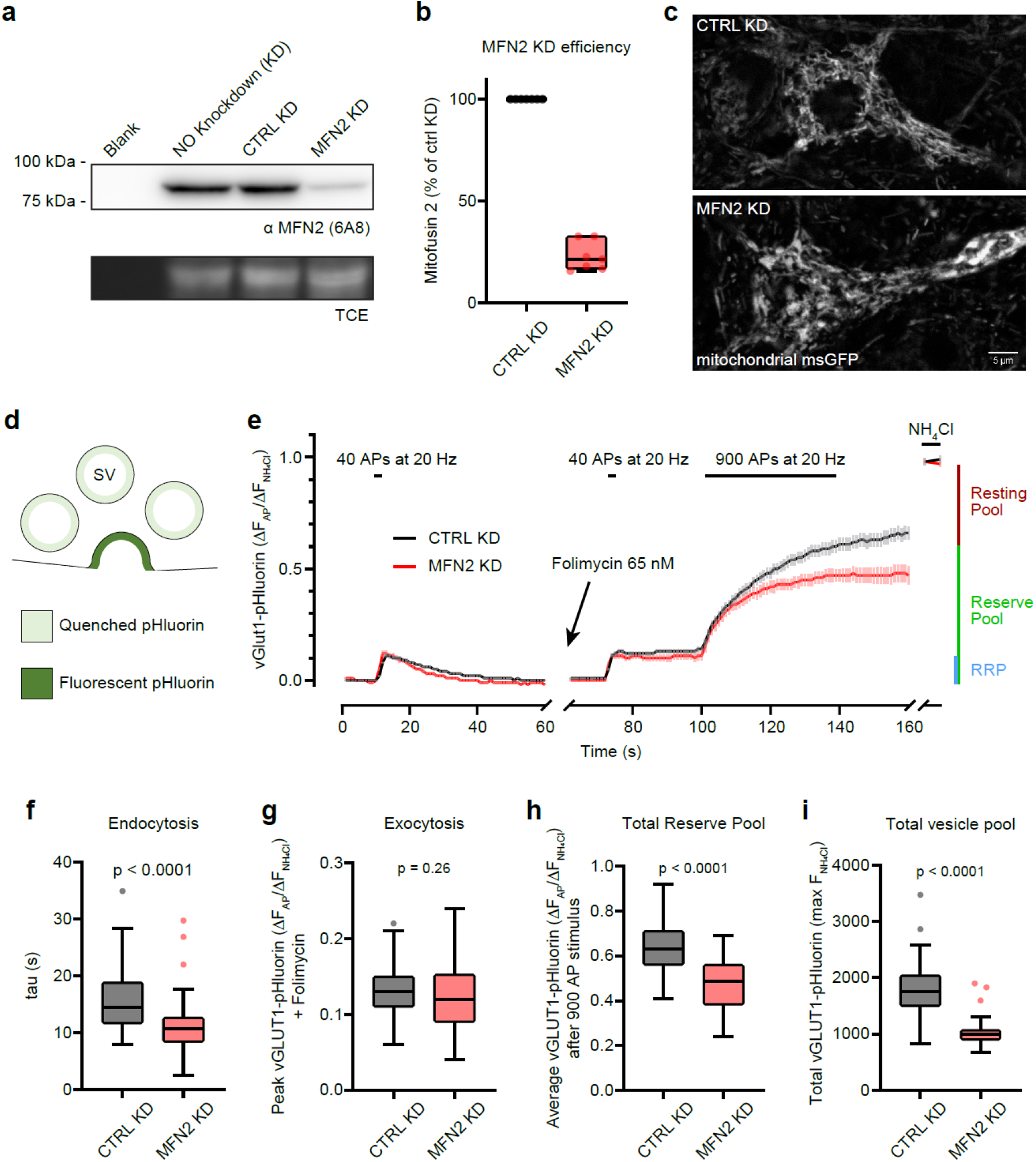
Mitochondrial fusion supports synaptic vesicle recycling. **a**) Representative anti-MFN2 immunoblot from rat 15 DIV hippocampal neurons with trichloroethanol (TCE) as a loading control. Conditions from left to right are blank/no protein, control – no lentivirus application, control – non-targeting shRNA lentivirus (CTRL KD), and sample treated with MFN2 targeted shRNA (see materials and methods) (MFNs KD). **b**) A Tukey box and whisker plot of MFN2 knockdown efficiency quantitated by immunoblot densitometry from 7 independent dissociated neuronal samples. Median MFN KD (red) percentage is 21.45% [95% CI 15.6% - 32.7%], Wilcoxon Signed Rank test, p = 0.0156. Detailed statistics in Figure 3 – source data 1. **c**) Representative Airyscan images of single focal planes through the soma and proximal dendrites of neurons transduced with mitochondrial-matrix targeted msGFP, and CTRL or MFN2 KD lentivirus constructs. Mitochondrial aggregation and loss of tubular network, characteristic of loss of MFN2 function, is visible in the MFN2 KD treated neurons. Scale bar: 5 µm. **d**) Schematic of SV targeted pHluorin during exocytosis. **e**) Average (+/- 95% CI) traces of vGlut1-pHluorin fluorescence intensity during the synaptic vesicle cycle from CTRL KD (black) and MFN2 KD (red) neurons. Neurons were stimulated with 40 action potentials (40 AP) at 20 Hz to drive the release of the readily releasable pool (RRP) and then allowed to recover. During recovery, the fluorescence decay represents endocytosis and vesicle reacidification. After recovery, folimycin was added to prevent vesicle reacidification (pHluorin quenching) and allow a measure of pure exocytosis. Forty AP were used again to drive the release of the RRP; the neurons were then maximally stimulated with 900 AP at 20 Hz to trigger exocytosis of the entire RP of vesicles. After stimulation, NH_4_Cl was perfused onto the sample to unquench all synaptic pHluorin signal and measure the total vGlut1-pHluorin signal. Traces normalized to maximum NH_4_Cl signal ((F – F_0_)/(F_max_ – F_0_)), n > 45 separate boutons from 3 independent experiments. **f**) Rate of endocytosis (τ) determined from the vGlut1-pHluorin fluorescence decay after 40 AP stimulation. The median value for CTRL KD (black) boutons is 14.39 s [95% CI 13.17 – 17.85] and 10.73 s [95% CI 9.58 – 11.94] for MFN2 KD (red) boutons, Mann-Whitney test p < 0.0001, detailed statistics in Figure 3 – source data 2. **g**) The RRP as determined from the vGlut1-pHluorin fluorescence peak after incubation with 65 nM folimycin and 40 AP stimulation. The median value for CTRL KD (black) boutons is 13% [95% CI 11% – 14%] and 12% [95% CI 10% – 13%] for MFN2 KD (red) boutons, Mann-Whitney test p = 0.26, detailed statistics in Figure 3 – source data 3. **h**) Total RP fraction assayed measuring the vGlut1-pHluorin fluorescence peak after incubation with 65 nM folimycin and stimulation with 940 AP. The median value for CTRL KD (black) boutons is 63% [95% CI 59% – 67%] and 49% [95% CI 41% – 55%] for MFN2 KD (red) boutons, Mann-Whitney test p < 0.0001, detailed statistics in Figure 3 – source data 4. **i**) A measure of the total vesicle pool as assayed by vGlut1-pHluorin fluorescence during NH_4_Cl perfusion. The median value for CTRL KD (black) boutons is 1755 arbitrary fluorescence units (AFU) [95% CI 1642 - 1954] and 1002 AFU [95% CI 952 – 1018] for MFN2 KD (red) boutons, Mann-Whitney test p < 0.0001, detailed statistics in Figure 3 – source data 5. Charts (3f, 3g, 3h, and 3i) are Tukey box and whisker plots, quantified from acquired traces in (3e).

**Figure 3 Figure supplement 1.**
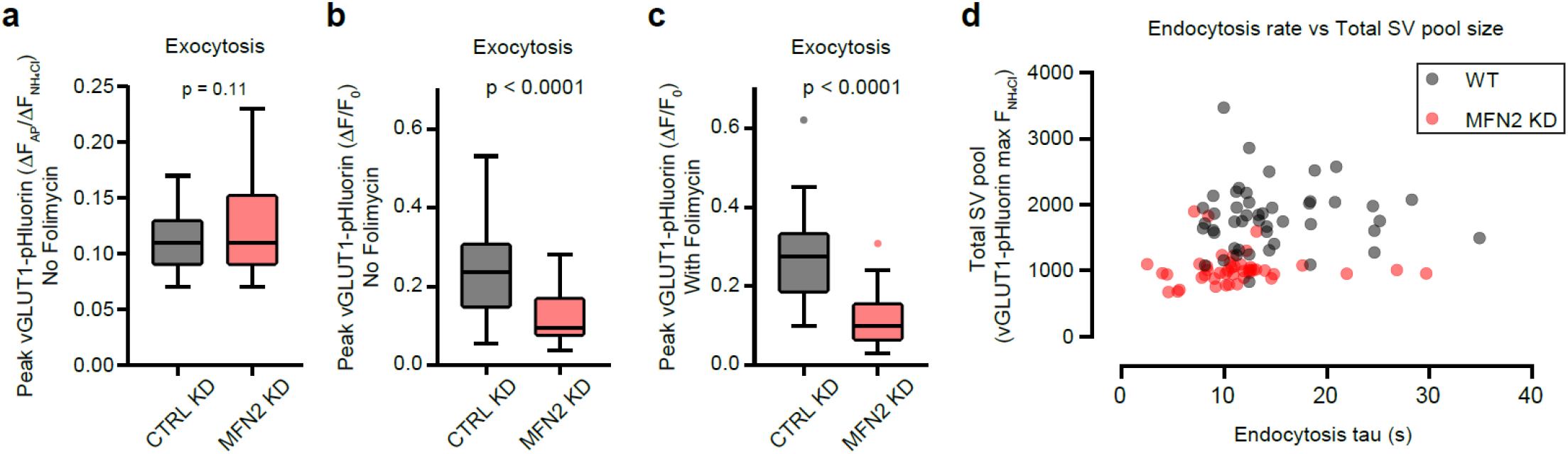
**a**) The RRP fraction as assayed by vGlut1-pHluorin fluorescence peak after 40 AP stimulation (no folimycin). The median value for CTRL KD boutons is 11% [95% CI 10% – 12%] and 11% [95% CI 10% – 15%] for MFN2 KD boutons, Mann-Whitney test p = 0.11, detailed statistics in Figure 3 Figure supplement 1 – source data 1. **b**) The RRP fraction as assayed by vGlut1-pHluorin fluorescence peak after 40 AP stimulation, but without folimycin and without NH_4_Cl normalization (compare to Figure 3 Figure supplement 1a). The median value for CTRL KD boutons is 24% [95% CI 17% – 27%] and 9% [95% CI 8% – 12%] for MFN2 KD boutons, Mann-Whitney test p < 0.0001, detailed statistics in Figure 3 Figure supplement 1 – source data 2. **c**) The RRP fraction as assayed by vGlut1-pHluorin fluorescence peak after 40 AP stimulation, with folimycin and without NH_4_Cl normalization (compare to Figure 3g). The median value for CTRL KD boutons is 24% [95% CI 17% – 27%] and 9% [95% CI 8% – 12%] for MFN2 KD boutons, Mann-Whitney test p < 0.0001, detailed statistics in Figure 3 Figure supplement 1 – source data 3. **d**) An X-Y scatterplot of total vesicle pool values against endocytosis values calculated in (3f) and (3i). MFN2 KD boutons are generally smaller and have faster rates of endocytosis than their CTRL KD counterparts. Charts (1a, 1b and 1c) are Tukey box and whisker plots, quantified from acquired traces in (3e).

### Mitochondrial fusion supports calcium homeostasis and the synaptic proteome

To gain insight into our pHluorin measurements (Figure 3), we next investigated presynaptic Ca^2+^ ([Ca^2+^]_i_) using synaptic targeted HTL-JF646-BAPTA-AM **(Bradberry & Chapman, 2022; Deo et al., 2019)**. With a stimulation paradigm of a single AP, followed by a short recovery and a sensor-saturating stimulation of 50 AP at 50 Hz, followed finally by a longer recovery; we calculated resting Ca^2+^, Ca^2+^ entry following 1 AP, as well as Ca^2+^ decay from 1 AP or after a train. Average traces from CTRL and MFN2 KD presynapses are shown in Figure 4a. We find that there is a trend toward lower resting Ca^2+^ in the MFN2 KD condition (Figure 4b) and no difference in Ca^2+^ entry in response to 1 AP (Figure 4c). However, Ca^2+^ decay in the MFN2 KD condition, was altered. Ca^2+^ decay following 1 AP was well fitted by a single-exponential function, while decay from 50 AP required a double-exponential function. These observations agree well with previous studies that find longer bursts of activity elicit a Ca^2+^ decay that is well fitted to a double-exponential function **(Koester & Sakmann, 2000; Tank, Regehr, & Delaney, 1995; Zhang & Linden, 2012)**, with the second component mediated by mitochondrial Ca^2+^ flux **(Werth & Thayer, 1994)**. Following a single AP, [Ca^2+^]_i_ decreases much faster in the MFN2 KD condition versus the CTRL KD condition (Figure 4d), with the best fit exponentials compared in (Figure 4e). Following a 50 AP train, both time constants (τ) representing [Ca^2+^]_i_ decay are also faster in the MFN2 KD condition (Figure 4f), with the best fit exponentials compared in (Figure 4g). The τ from 1 AP is the same as the [FAST] τ from the train, suggesting that the same [Ca^2+^]_i_ decay mechanism governs both processes (Figure 4h).

**Figure 4.**
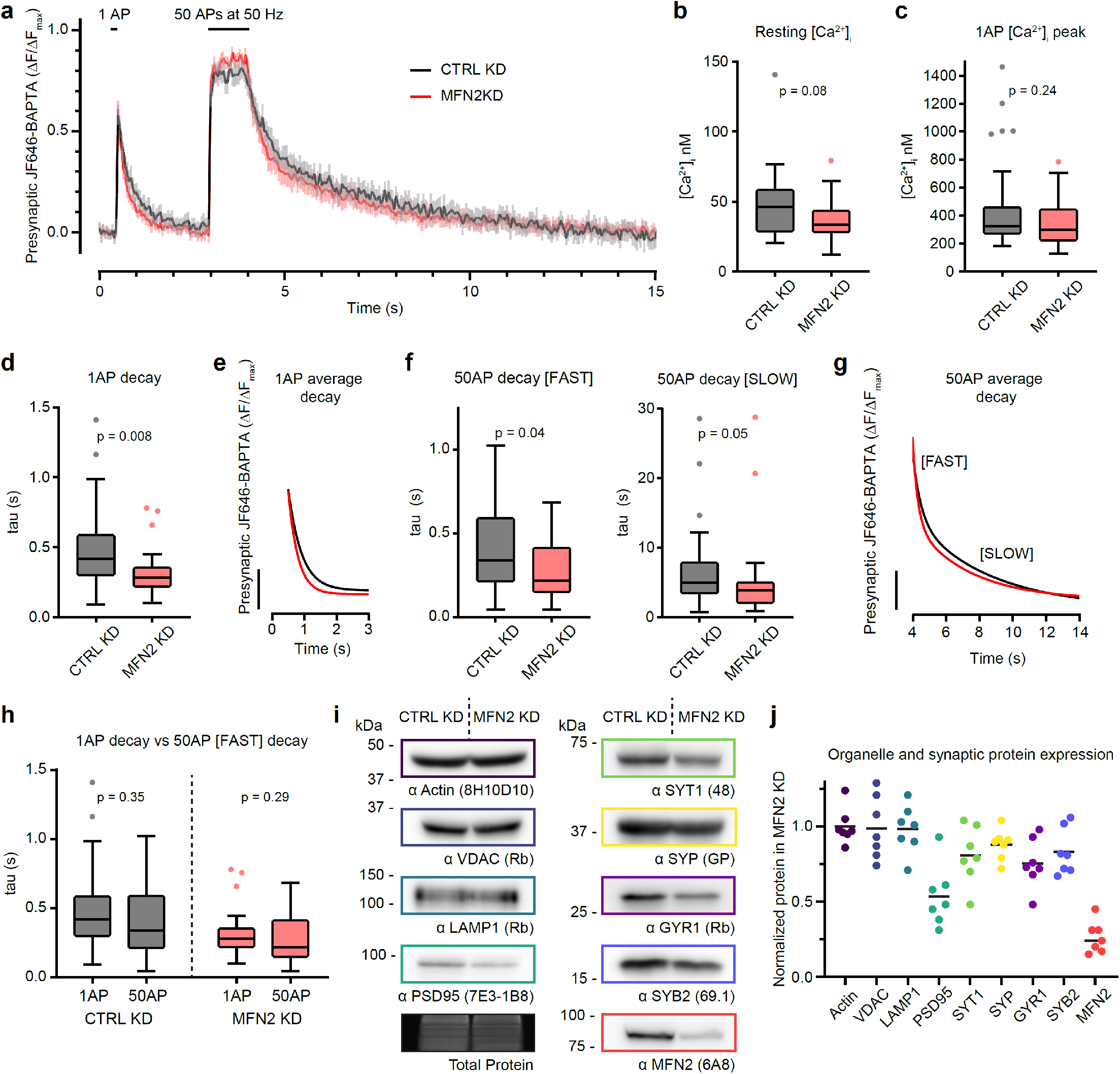
Mitochondrial fusion supports presynaptic calcium dynamics and the synaptic proteome. **a**) Average (+/- 95% CI) traces of synaptophysin-HaloTag (SYP-HT) bound HTL-JF646-BAPTA fluorescence intensity during electrical stimulation from CTRL KD (black) and MFN2 KD (red) neurons. Neurons were stimulated with a single action potential (1 AP) to quantitate calcium (Ca2^+^) influx. After 2.5 seconds, the neurons were maximally stimulated by 50 AP at 50 Hz to saturate the HTL-JF646-BAPTA and calculate absolute Ca2^+^ concentration calculations as described in Materials and Methods. Traces are from n = 30 separate boutons, from 3 independent experiments. **b**) Resting presynaptic Ca^2+^ ([Ca^2+^]_i_) from CTRL KD (black) and MFN2 KD (red) boutons, median values were 46.2 nM [95% CI 32 - 54] and 33.4 nM [95% CI 29 – 37], respectively. Mann-Whitney test p = 0.0765, detailed statistics in Figure 4 – source data 1. **c**) Single action potential peak [Ca^2+^]_i_ from CTRL KD (black) and MFN2 KD (red) boutons, median values were 325.5 nM [95% CI 282 - 380] and 298.1 nM [95% CI 235 – 381.4], respectively. Mann-Whitney test p = 0.2403, detailed statistics in Figure 4 – source data 2. **d**) Single action potential [Ca^2+^]_i_ τ from CTRL KD (black) and MFN2 KD (red) boutons. Average traces were well fitted to a single-exponential function, and median values were 419 ms [95% CI 372 - 540] and 283 ms [95% CI 242 – 312], for CTRL KD and MFN2 KD, respectively. Mann-Whitney test p = 0.0081, detailed statistics in Figure 4 – source data 3. **e**) Single-exponential best fit curves to average CTRL KD (black) and MFN2 KD (red) [Ca^2+^]_i_ decay traces from (4a), detailed values in Figure 4 – source data 4. **f**) Train action potential (50 AP at 50 Hz) [Ca^2+^]_i_ τ from CTRL KD (black) and MFN2 KD (red) boutons. Average traces well fitted with a double-exponential function, and median values for the [FAST] component were 339 ms [95% CI 250 - 513] and 218 ms [95% CI 165 – 361], for CTRL KD (black) and MFN2 KD (red) respectively, Mann-Whitney test p = 0.0417. Median values for the [SLOW] component were 4.9 s [95% CI 3.5 – 7.1] and 3.8 s [95% CI 2.3 – 4.5], for CTRL KD (black) and MFN2 KD (red) respectively, Mann-Whitney test p = 0.0489, detailed statistics in Figure 4 – source data 5. **g**) Double-exponential best fit curves for the average CTRL KD (black) and MFN2 KD (red) [Ca^2+^]_i_ decay traces from (4a), detailed values are in Figure 4 – source data 6. **h**) Comparison of CTRL KD (black) and MFN2 KD (red) τ from (4d) and the [FAST] component from (4f). The [FAST] component is not significantly different from the single component from 1 AP, Mann-Whitney test p = 0.3455 and p = 0.2942 for CTRL KD (black) and MFN2 KD (red), respectively. Detailed values are in Figure 4 – source data 7. Charts (4b, 4c, 4d, 4f, and 4h) are Tukey box and whisker plots, quantified from acquired traces in (4a). **i**) Representative immunoblots used to quantify the indicated synaptic and organelle protein levels in CTRL KD and MFN2 KD neurons. Protein detected and antibody (clone if monoclonal, species if polyclonal) are listed below the blots, and total protein was assayed using Lumitein (Biotium) for every condition, including repeated lanes. **j**) Quantitation of protein levels in MFN2 KD samples, normalized to CTRL KD and total protein. Samples are n = 7 from 7 independent trials, detailed values are in Figure 4 – source data 8.

Finally, we followed up on the result from Figure 3i, where we measured a decreased amount of transduced pHluorin reporter reaching the bouton, suggesting that SVs were decreased by approximately half. We compared protein levels of housekeeping (Actin) and general organelle (voltage dependent anion channel, VDAC; lysosome-associated membrane protein 1, LAMP1) markers to presynaptic (synaptotagmin 1, SYT1; synaptophysin, SYP; synaptogyrin 1, GYR1; synaptobrevin 2, SYB2) and a postsynaptic marker (post-synaptic density 95, PSD95) between CTRL and MFN2 KD neurons (Figure 4i). Protein levels of Actin, VDAC, and LAMP1 are unaltered in MFN2 KD neurons, while presynaptic protein markers are decreased by ∼25%, and PSD95 is decreased by ∼50%. In these trials, MFN2 was decreased by ∼75% (Figure 4j).

These experiments revealed that [Ca^2+^]_i_ decay and SV protein homeostasis is disrupted when mitochondrial fusion is compromised. An altered SV proteome and faster Ca^2+^ transients can explain SV cycle defects demonstrated by our pHluorin experiments, which we discuss below.

## Discussion

### Mitochondrial trafficking asymmetry in axons

To address the existence of axonal mitochondrial QC pathways, analysis of mitochondrial morphology, trafficking, and function in axons after application of high, moderate, or low doses of mitochondrial specific toxins (mito-toxins), have been conducted. Primarily, these toxins target either the mitochondrial membrane potential (ΔΨ_m_) or components of the electron transport chain. High doses of Antimycin A (AA, >40 µM), a complex III inhibitor, either arrested mitochondrial transport **(Wang et al., 2011)** and induced axonal mitophagy **(Ashrafi et al., 2014)**; or increased axonal retrograde flux of mitochondria **(Miller & Sheetz, 2004)**. A high dose of Paraquat (20 mM) specifically inhibits mitochondrial traffic in the axon, dependent on reactive oxygen species (ROS) **(Liao, Tandarich, & Hollenbeck, 2017)**. A moderate dose of carbonyl cyanide m-chlorophenyl hydrazone (CCCP, 10 µM), a ΔΨ_m_ dissipating reagent, increased axonal retrograde traffic to the soma **(Cai et al., 2012)**, or at a high dose, 1 mM, had no effect on the retrograde flux of mitochondria **(Miller & Sheetz, 2004)**. Low doses of AA (<10 nM) increased retrograde mitochondrial transport **(M. Y. Lin et al., 2017)**, or had no effect on motility but induced mitophagy in the soma **(Evans & Holzbaur, 2020)**. During basal or unstressed conditions, anterograde and retrograde trafficked mitochondria are reported to either correlate strongly **(Miller & Sheetz, 2004)**, or to not correlate at all **(Verburg & Hollenbeck, 2008)**, with ΔΨ_m_, or ATP levels **(Suzuki et al., 2018)**; however, inhibition of retrograde axonal traffic increased the number of damaged mitochondria in axons **(Mandal et al., 2021)**.

Here, we document differences in mitochondrial trafficking between axons and dendrites, as well as trafficking asymmetry within axons. Anterograde axonal mitochondria are larger and relatively more reduced, while retrograde axonal mitochondria are shorter and relatively more oxidized (Figure 1). Importantly, our studies are done in mature, unstressed primary dissociated hippocampal cultures, and we use microfluidics and MitoTracker dyes to unambiguously define anterograde and retrograde axonal mitochondria; the mitochondria we observe in the middle of the microchannel have trafficked a minimum of 150 microns in a single direction. We believe this is the primary reason we see such a clear distinction between anterograde and retrograde populations. Importantly, we used MitoTracker Red as an axonal mitochondrial stain and therefore likely over-estimate the size of retrograde trafficked mitochondria, as only the mitochondria with ΔΨ_m_ were detected. In sum, our data demonstrates a trafficking asymmetry of mitochondria in the axon, which we interpret as evidence of an active QC mechanism.

### Role of mitochondrial fusion in maintaining the axonal mitochondrial network

Anterograde mitochondria that are larger and more fit could serve to either establish new mitochondrial outposts or replenish (complement) existing, stationary axonal mitochondria. In our studies, we document axonal mitochondrial complementation and find that total mitochondrial complementation occurs at a rate of approximately 2% per hour (Figure 2). Importantly, approximately 50% of new axonal mitochondria (MTG+) fuse with existing axonal mitochondria over the course of our experiments. This means that half of the newly trafficked mitochondria serve to immediately complement existing axonal mitochondria. The fate of the other half of the newly trafficked mitochondria is unclear, as these may be establishing new axonal outposts, or simply continuing their journey down the axon for future complementation events. These experiments demonstrate the dynamic fusion properties of mitochondria in the axon and suggest that continual mitochondrial complementation may be important to support the mitochondrial network and axon function (i.e., exocytosis). Indeed, mitochondrial complementation (via fusion) suppresses mtDNA disease associated alleles in various mouse tissues **(Nakada et al., 2001)**, and loss of mitochondrial fusion increases mitochondrial heterogeneity and dysfunctional mitochondrial units in mouse embryonic fibroblasts (MEFs) **(Chen, Chomyn, & Chan, 2005)**. In INS1 and COS7 cells, fusion is selective and precedes fission events **(Twig et al., 2008)** and in rat dorsal root ganglion (DRG) neurons, fusion hyperpolarizes mitochondria and promotes ATP synthesis **(Suzuki et al., 2018)**.

### Mitochondrial fusion supports the synaptic vesicle cycle

We demonstrated that there is a mitochondrial trafficking asymmetry in axons and that mitochondrial fusion occurs frequently and can complement stationary mitochondria. We then investigated the role of mitochondrial fusion in supporting the main function of the axon, the SV cycle, by knocking down MFN2 and expressing SV targeted pHluorin. Altering the expression of mitochondrial fusion or fission machinery will alter the morphology of mitochondria in the axon **(Amiri & Hollenbeck, 2008)**. Disrupting mitochondrial fission leads to loss of mitochondria in dopaminergic axons **(Berthet et al., 2014)**, decreased neurite and synapse formation in primary neuronal cultures **(Ishihara et al., 2009)**, and enhanced depression in acute hippocampal slices **(Oettinghaus et al., 2016)**. Disrupting mutations in MFN2 infamously drive a progressive form of Charcot-Marie-Tooth syndrome (type 2A) and alter axonal mitochondrial localization **(Baloh, Schmidt, Pestronk, & Milbrandt, 2007; A. Misko, Jiang, Wegorzewska, Milbrandt, & Baloh, 2010; A. L. Misko, Sasaki, Tuck, Milbrandt, & Baloh, 2012)**. Suppressing MFN2 expression in human-induced pluripotent stem cell (hiPSC) derived neurons inhibits differentiation and synaptogenesis **(Fang, Yan, Yu, Chen, & Yan, 2016)**. We find that disruption of mitochondrial fusion by MFN2 KD leads to the decreased expression of SV-associated proteins, decreased absolute exocytosis, difficulty in mobilizing the reserve pool of SVs, and faster endocytosis (Figure 3).

Here we focused on disrupting fusion, but disrupting fission also inhibits the mobilization of the reserve pool at *Drosophila* neuromuscular junctions (NMJ) **(Verstreken et al., 2005)**, suggesting that mitochondrial localization or morphology influence the SV cycle. However, additional studies in *Drosophila* suggest that while mitochondrial morphology is critical for localization and trafficking in the axon, it is dispensable for normal function of the neuron **(Trevisan et al., 2018)**. Moreover, non-functional mitochondria fail to buffer presynaptic Ca^2+^ and increase synaptic depression rates; this depression is completely rescued by supplementing the buffering of cytosolic Ca^2+^ **(Billups & Forsythe, 2002)**. Altering the size of mitochondria changes their Ca^2+^ buffering capacity. Smaller mitochondria in MFN2 KD C2C12 myotubes buffer less Ca^2+^ and have lower resting [Ca^2+^]_i_ **(Kowaltowski et al., 2019)**, but see also **(Naon et al., 2016)**. Disrupting fission in C2C12 myotubes **(Kowaltowski et al., 2019)** or in mouse pyramidal neurons **(Lewis et al., 2018)** enhances mitochondrial Ca^2+^ buffering capacity. Generally, mitochondria are shown to influence presynaptic Ca^2+^ transients and in turn, modulate presynaptic properties [recently reviewed in **(Datta & Jaiswal, 2021)**]. Our pHluorin results prompted us to investigate presynaptic Ca^2+^ after MFN2 KD.

The recently developed far-red BAPTA based indicator JF646-BAPTA-AM **(Deo et al., 2019)** allowed us to calculate (see Materials and Methods) resting presynaptic Ca^2+^ ([Ca^2+^]_i_), Ca^2+^ influx, and [Ca^2+^]_i_ decay kinetics. The rate of [Ca^2+^]_i_ decay measured here is not reflective of the absolute efflux rate because we use a high-affinity indicator **(Regehr & Atluri, 1995)**, but relative differences between conditions are still informative. We found a trend toward decreased resting [Ca^2+^]_i_ and no difference in Ca^2+^ influx from a single AP when comparing CTRL to MFN2 KD neurons. We also found that Ca^2+^ decay was faster after a single AP or a train in MFN2 KD neurons (Figure 4). Mitochondria are a key Ca^2+^ buffer at the presynapse and loss of this mitochondrial function would speed presynaptic Ca^2+^ transients **(Werth & Thayer, 1994)**. Indeed, loss of the mitochondrial Ca^2+^ uniporter (MCU) increases the rate of SV endocytosis in mouse hippocampal neurons **(Marland, Hasel, Bonnycastle, & Cousin, 2016)**, however the role of Ca^2+^ in endocytosis is complex. Increased [Ca^2+^]_i_ either promotes **(Balaji, Armbruster, & Ryan, 2008; Neher & Zucker, 1993; Neves, Gomis, & Lagnado, 2001; Sankaranarayanan & Ryan, 2001; Wu, Xu, Wu, & Wu, 2005)** or inhibits endocytosis **(Leitz & Kavalali, 2011; Sankaranarayanan & Ryan, 2000; von Gersdorff & Matthews, 1994)**, see also a thorough review **(Leitz & Kavalali, 2016)**. Increased levels of exocytosis can also saturate the endocytotic machinery and prolong the rate of endocytosis **(Armbruster, Messa, Ferguson, De Camilli, & Ryan, 2013; Delvendahl, Vyleta, von Gersdorff, & Hallermann, 2016; Sankaranarayanan & Ryan, 2000; Sun, Wu, & Wu, 2002)**. When we KD MFN2, we observed reductions in the extent of exocytosis and faster Ca^2+^ transients. Therefore, even though it may be counterintuitive, it stands to reason that the observed endocytosis rate was faster in the MFN2 KD.

### Final thoughts

MFN2 is a large membrane protein with numerous cellular functions. Its original and most widely recognized function is to catalyze the homotypic fusion of outer mitochondrial membranes **(Chen et al., 2003)**. It also either positively **(de Brito & Scorrano, 2008; Naon et al., 2016)** or negatively **(Cosson, Marchetti, Ravazzola, & Orci, 2012; Filadi et al., 2015)** regulates mitochondria-ER contact sites and may interact with motor adaptors in vertebrates **(A. Misko et al., 2010; A. L. Misko et al., 2012)**. Therefore, the interpretation of phenotypes resulting from constitutive knockouts or even acute KD experiments, as done here, should be made with caution. Regardless, these experiments are essential to begin to understand the role of mitochondrial complementation (fusion) in supporting the axonal mitochondrial network and the physiology of the axon. More acute methods, to disrupt this membrane bound protein, will further clarify its role in presynaptic physiology. Additionally, it will also be important to selectively disrupt motifs or domains in MFN2. Acute and domain specific disruption of function can be implemented with a tool like knockoff **(Vevea & Chapman, 2020)**. The topology of MFN2 is also debated, it is anchored either by a double **(Rojo, Legros, Chateau, & Lombes, 2002)** or single **(Mattie, Riemer, Wideman, & McBride, 2018)** transmembrane domain. Experiments based on the knockoff approach could shed additional light on this topology. Moreover, a surprising result from our experiments is the stark decreased expression of PSD95 in MFN2 KD neurons. An interesting question is whether this effect is mediated by pre- or postsynaptic mitochondria; compartment specific knockoff could be used to address this issue.

Exciting questions remain as to the fundamental signals that govern the maintenance and QC of the axonal mitochondrial network. A large body of research revealed that loss of ΔΨ_m_ marks a mitochondrion for degredation, and it is unable to rejoin the network **(Pickles, Vigie, & Youle, 2018)**, but what is the signal that identifies a mitochondrion for a complementation event? How are mitochondrial fusion and fission events linked? MFN2 itself is sensitive to ROS and may provide an elegant mechanism to preferentially activate the fusion machinery **(Mattie et al., 2018; Shutt, Geoffrion, Milne, & McBride, 2012)**. Furthermore, how are mitochondria selected for axonal versus dendritic egress from the soma? Is there a QC step at this early stage in trafficking? What are some of the damage markers that build up over time, and can they be influenced? To address these questions, methods that facilitate the rapid isolation of naturally damaged or aged mitochondria from the axon need to be developed. These questions are important for understanding mitochondrial quality control mechanisms and how they support the SV cycle and neuronal health.

## Acknowledgements

We would like to thank the Chapman lab members for valuable discussions related to this manuscript and C. Greer and E.T. Watson specifically for critical reading of the manuscript and valuable edits. We would also like to thank D.T. Larson, J.H. Rinald and B. Stumpner for excellent technical assistance. This study was supported by grants from the NIH (MH061876 and NS097362 to E.R.C). J.D.V. was supported by a postdoctoral fellowship from the NIH F32 NS098604 and the Warren Alpert Distinguished Scholars Fellowship award. E.R.C. is an Investigator of the Howard Hughes Medical Institute.

## Declaration of Interests

The authors declare no competing financial interests.

## Methods

| Key Resource Table |  |  |  |  |
| --- | --- | --- | --- | --- |
| Reagent type (species) or resource | Designation | Source or reference | Identifiers | Additional information |
| biological sample (*Rattus norvegicus*) | Primary rat hippocampal neurons | Envigo | Sprague Dawley |  |
| cell line (*Homo-sapiens*) | HEK293T, Kidney epithelial | ATCC | CRL-11268 |  |
| recombinant DNA reagent | FUGW (plasmid) | addgene | Addgene plasmid # 14883; http://n2t.net/addgene:14883 ; RRID:Addgene_14883 | Lentivirus backbone |
| recombinant DNA reagent | pEF-GFP (plasmid) | addgene | Addgene plasmid # 11154; http://n2t.net/addgene:11154 ; RRID:Addgene_11154 | pEF backbone |
| recombinant DNA reagent | pEF cyto-cmCh (plasmid) | This study | Addgene | Materials and methods |
| recombinant DNA reagent | pEF mito-cGFP (matrix) (plasmid) | This study | Addgene | Materials and methods |
| recombinant DNA reagent | pEF mito-PAcGFP (plasmid) | This study | Addgene | Materials and methods |
| recombinant DNA reagent | pF(UG) hSyn vGlut1 pHluorin (plasmid) | This study | Addgene | Materials and methods |
| recombinant DNA reagent | pEGFP-N1-mt-ro1GFP (plasmid) | Addgene | Addgene plasmid # 82407 ; http://n2t.net/addgene:82407 ; RRID:Addgene_82407 |  |
| recombinant DNA reagent | TRC2 pLKO.5-puro Non-Mammalian shRNA Control (plasmid) | Sigma | SHC202 | CTRL KD |
| recombinant DNA reagent | pLKO.1 MFN2KD (plasmid) | Sigma | TRCN0000080612 | MFN2 KD |
| recombinant DNA reagent | pF(UG) hSyn SYP-HaloTag (plasmid) | This study | addgene | Materials and methods |
| Antibody | anti-MFN2 (mouse monoclonal) | Abcam; Abnova | Abcam Cat# ab56889, RRID:AB_2142629; Abnova Cat# H00009927-M01, RRID:AB_714775 | IB (1:1000) |
| Antibody | anti-SYT1 (mouse monoclonal) | DSHB | mAB 48; RRID:AB_2199314 | IB (1:1000)<br>ICC (1:100) |
| Antibody | anti-SYP (guinea pig polyclonal) | SySy | 101 004; RRID:AB_1210382 | IB (1:1000)<br>ICC (1:500) |
| Antibody | anti-VDAC (Rabbit polyclonal) | EMD/ Sigma-Aldrich | AB10527 \| Anti-VDAC Antibody; RRID:AB_10806766 | IB (1:1000) |
| Antibody | anti-LAMP1 (Rabbit polyclonal) | Abcam | ab24170; RRID:AB_775978 | IB (1:500) |
| Antibody | anti-PSD95 (mouse monoclonal) | ThermoFisher | Clone 7E3-1B8; Catalog # MA1-046; RRID:AB_2092361 | IB (1:1000) |
| Antibody | anti-SYB2 (mouse monoclonal) | SySy | Clone 69.1; catalog #104 211C5, RRID:AB_2619757 | IB (1:1000) |
| Antibody | anti-GYR1 (polyclonal rabbit) | SySy | 103 002, RRID:AB_887818 | IB (1:1000) |
| Antibody | goat anti-guinea pig IgG-HRP | Abcam | ab6908, RRID:AB_955425 | IB (1:10,000) |
| Antibody | goat anti-rabbit IgG-HRP | Biorad | 1706515, RRID:AB_11125142 | IB (1:10,000) |
| Antibody | goat anti-mouse IgG2b-HRP | Biorad | M32407; RRID:AB_2536647 | IB (1:10,000) |
| Antibody | goat anti-mouse IgG-HRP | Biorad | 1706516; RRID:AB_11125547 | IB (1:10,000) |
| Chemical | MitoTracker Green FM | ThermoFisher | M7514 |  |
| Chemical | MitoTracker Red CMXRos | ThermoFisher | M7512 |  |
| Chemical | Folimycin / Concanamycin A | Tocris | 2656 |  |
| Chemical | Trichloroethanol (TCE) | Sigma | T54801-100G | Total protein stain |
| Material | Xona microfluidic | Xona | SND450 | Standard neuron microfluidic |
| Chemical | HTL-JF646-BAPTA-AM | Janelia Research Campus/ HHMI | A kind gift from Dr. Luke Lavis | Luke Lavis Lab (Deo et al., 2019) |

### Ethics Statement

Animal care and use in this study were conducted under guidelines set by the NIH Guide for the Care and Use of Laboratory Animals handbook. Protocols reviewed and approved by the Animal Care and Use Committee (ACUC) at the University of Wisconsin, Madison (Laboratory Animal Welfare Public Health Service Assurance Number: A3368-01).

### Cell Culture

Rat (Sprague Dawley) hippocampal and cortical neurons were isolated from E18 pups (Envigo) using a procedure previously described in **(Vevea & Chapman, 2020)**. In brief, rat hippocampal neurons were dissected, trypsinized (Corning; 25-053-CI), triturated, and plated on glass coverslips (Warner instruments; 64-0734 (CS-18R17)), previously coated with poly-D-lysine (Thermofisher; ICN10269491). Hippocampal neurons were also cultured in standard neuron microfluidic devices (SND450, XONA Microfluidics) mounted on glass coverslips, as previously described **(Bomba-Warczak et al., 2016)**. Neuronal cultures were grown in Neurobasal-A (Thermofisher; 10888-022) medium supplemented with B-27 (2% Thermofisher; 17504001), Glutamax (2 mM Gibco; 35050061), and pen/strep before experiments. All experiments were performed between 13 to 20 days in vitro (DIV); 13-16 DIV for microfluidic experiments and 15-20 DIV for pHluorin and Calcium imaging. For lentivirus preparation, HEK293T cells (ATCC) were cultured following ATCC guidelines. These cells had been previously tested for mycoplasma contamination using the Universal Mycoplasma Detection Kit (ATCC; 30-1012K). The HEK293T cells were also validated using Short Tandem Repeat profiling by ATCC (ATCC; 135-XV).

### Lentivirus production and use

Lentivirus production was performed as described previously **(Vevea & Chapman, 2020)**. When needed, lentiviral constructs were subcloned into the FUGW transfer plasmid (FUGW was a gift from David Baltimore (Addgene plasmid # 14883 ; http://n2t.net/addgene:14883 ; RRID:Addgene_14883)) **(Lois, Hong, Pease, Brown, & Baltimore, 2002)**. We had previously replaced the ubiquitin promoter with the CAMKII promoter or human synapsin I promoter **(Kugler, Kilic, & Bahr, 2003; Vevea & Chapman, 2020)**. Lentivirus was added to neuronal cultures between 2 DIV and 5 DIV.

### Plasmid use and construction

The redox sensitive green fluorescent protein (pEGFP-N1-mt-ro1GFP) was a gift from S. James Remington (Addgene plasmid # 82407 ; http://n2t.net/addgene:82407 ; RRID:Addgene_82407). Silencing constructs encoding control shRNA and *mfn2* shRNA (target sequence: TGGATGGACTATGCTAGTGAA) were purchased from Sigma (SHC202 and TRCN0000080612 respectively). The vGLUT1-pHluorin construct **(Voglmaier et al., 2006)** was subcloned into our previously modified lentivirus backbone of choice derived from FUGW **(Vevea & Chapman, 2020)**. For measuring synaptic Ca^2+^, the HaloTag cassette from pHTC HaloTag CMV-neo Vector (Promega; G7711), was PCR-amplified and appended to the carboxy-terminus of synaptophysin (SYP) with a GS(GSS)_4_ linker, and subcloned into the modified FUGW lentiviral vector. Mitochondrial matrix targeted GFP, cytosolic mCherry, and photoactivatable GFP were subcloned into pEF-GFP after excising the GFP (pEF-GFP was a gift from Connie Cepko (Addgene plasmid # 11154 ; http://n2t.net/addgene:11154 ; RRID:Addgene_11154)) **(Matsuda & Cepko, 2004)**. Note the fluorescent proteins GFP, PA-GFP, and mCherry used here lack the carboxy terminal MDELYK sequence, as this was found to promote protein aggregation in long-lived cells (unpublished observation, Dr. Joseph S. Briguglio).

### Live-cell imaging

Primary rat hippocampal cultures were transiently transfected with pEF cyto-cmCh and pEF mito-cGFP (matrix) (Fig. 1a), pEF mito-PAcGFP (Fig. 1b), or pEGFP-N1-mt-ro1GFP (Fig. 2a-d) at 2-5 DIV using the Calcium Phosphate protocol **(Jordan & Wurm, 2004)**. For microfluidic experiments, neurons were transfected in the soma chamber, or labeled with MitoTracker Green FM (ThermoFisher) in the soma and MitoTracker Red CMXRos (Thermofisher) in the axon chamber. Microfluidic chambers (soma and axon) were sequentially labeled, being sure to keep unstained chamber pressure high during staining to maintain fluidic isolation. Other neuronal cultures were transduced with vGLUT1-pHluorin and knockdown (control and MFN2) lentiviruses at 2-5 DIV. Neuronal cultures were moved to standard imaging media (ECF (extracellular fluid/ECF) consisting of 140 mM NaCl, 5 mM KCl, 2 mM CaCl2, 2 mM MgCl2, 5.5 mM glucose, 20 mM HEPES (pH 7.3), B27 (Gibco), glutamax (Gibco), loaded into the microscope and maintained in a humidity-controlled chamber at near physiological temperature (35°C). Cultures were imaged using an Olympus FV1000 confocal microscope with a 60x 1.40 NA oil objective, using fixed laser intensity and gain settings (Fig. 1a) and on an Olympus IX83 inverted microscope equipped with a cellTIRF 4Line excitation system using an Olympus 60x/1.49 NA Apo N objective and an Orca Flash4.0 CMOS camera (Hamamatsu Photonics) running Metamorph software that was modified to run concurrently with Olympus 7.8.6.0 acquisition software from Molecular devices (Figure 1b-f; Figure 2; Figure 3e-I; Figure 4a-h). Photoactivation experiments were conducted using a single, spot focused, 405 nm laser on the CellTIRF system. Mitochondrial trafficking experiments were also conducted on the CellTIRF system using wide-field acquisition mode. Image sequences analyzing mitochondrial motility were acquired with five second intervals for a total of five minutes. Field electrical stimulation was triggered by a Grass SD9 stimulator run by Clampex 10.7.0.3 software (Molecular Devices) through a Digidata 1440A digitizer (Molecular Devices) to platinum parallel wires attached to a field stimulation chamber (Warner Instruments; RC-49MFSH). Stimulator voltage was set to 90v as this reliably elicited Ca^2+^ transients at every bouton. For pHluorin and calcium imaging experiments, D-AP5 (50 µM) (Abcam; ab120003), CNQX (20 µM) (Abcam; ab120044), Picrotoxin (100 µM) (Tocris; 1128) were added to the imaging media to prevent recurrent activity. **pHluorin imaging:** Synaptic vesicle exocytosis was monitored via change in fluorescence of presynaptic punctae from neurons transduced with the vGLUT1-pHluorin lentivirus. Images were acquired at 2×2 binning at 1 Hz for up to two minutes. For pHluorin imaging, the focal plane and field of view was found by live focusing during a test stimulation of 40 stimuli at 20 Hz. Neurons were given at least a minute to recover, and the stimulus was repeated during an image acquisition time of one minute. Next, freshly made folimycin (Tocris) 65 nM was added to the imaging well. The macrolide antibiotics bafilomycin and folimycin inhibit the function of vacuolar-type H+ -ATPase, preventing pHluorin quenching of recently endocytosed vesicles, allowing a measure of pure exocytosis **(Atasoy et al., 2008; Sankaranarayanan & Ryan, 2001)**. Within a minute of folimycin addition, a two-minute image acquisition was initiated and contained another pulse of 40 stimuli at 20 Hz, with a 30 second break, then a 900 stimuli train at 20 Hz **(Burrone, Li, & Murthy, 2006)**. At the end of this imaging sequence, a solution containing 50 mM NH_4_Cl (replacing 50 mM NaCl in ECF) was perfused onto the sample, dequenching all pHluorin **(Miesenbock, De Angelis, & Rothman, 1998). Calcium imaging:** Synaptic Ca^2+^ transients were recorded as synaptic fluorescence change from neurons transduced with synaptophysin-HaloTag (SYP-HT) reacted with HTL-JF646-BAPTA-AM (a gift from Dr Luke Lavis, Janelia Research Campus) **(Deo et al., 2019)**. Images were acquired at 2×2 binning at 20 Hz (50 ms exposure) for up to 20 seconds. Neurons were field stimulated with a single stimulus with a break for 2.5 seconds, then a 50-stimulus train at 50 Hz was given to saturate the indicator.

### Image Quantification

Mitochondrial motility (Fig. 1 and Fig. 2) was analyzed using Imaris 8.3 (Bitplane). Motile mitochondria were defined as having at least 3 (15 sec) consecutive displacements during image series. Mitochondrial redox ratio was calculated using FIJI (NIH) as described previously **(Vevea, Alessi Wolken, Swayne, White, & Pon, 2013)**. The roGFP redox ratio was normalized to the average ratio of each timeseries for main figures, non-normalized ratios are included in the supplement. pHluorin analysis was done by selecting responding ROIs in FIJI and measuring the fluorescence intensity change over time. These data were copied and imported into Axograph X 1.7.2 (Axograph Scientific) where traces were baseline subtracted and normalized to pHluorin peak fluorescence intensity during NH_4_Cl perfusion, ((F – F_0_)/(F_max_ – F_0_)). These traces were then analyzed in Axograph for endocytic τ (pHluorin signal decay after stimulus without folimycin), peak changes in fluorescence (exocytosis, with and without folimycin), and average fluorescence after 900 stimulus train but before NH_4_Cl application, which represents the total synaptic vesicle reserve pool (with folimycin). Calcium image quantitation was handled similarly to pHluorin analysis. Responding ROIs were identified, and fluorescence intensity was measured over time using FIJI. In axograph, traces were baseline subtracted and normalized to signal during the 50 AP train, ((F – F_0_)/(F_max_ – F_0_)). The decays of [Ca^2+^]_i_ after stimuli were measured as τ. The fluorescence decay after a 1 AP was well fit to a single-component exponential, while the decay after the train was well fit to a two-component exponential. Absolute [Ca^2+^]_i_ was quantitated as described previously **(Bradberry & Chapman, 2022)**. Briefly, the sensor was saturated (F_max_) with intense stimuli **(Maravall, Mainen, Sabatini, & Svoboda, 2000)**, and published reports of K_d_ (140 nM), R_f_ (5.5), and Hill coefficient (n) (1) **(Deo et al., 2019)**, were used to approximate [Ca^2+^]_i_ according to equations derived in **(de Juan-Sanz et al., 2017; Maravall et al., 2000)**. The exact following equation was used to calculate calcium concentration at the presynapse:

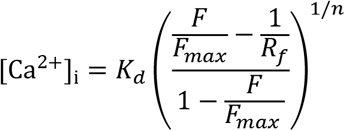

Where [Ca^2+^]_i_ represents internal calcium concentration, F is resting fluorescence, F_max_ is maximal fluorescence during indicator saturation, R_f_ is dynamic range, K_d_ is the dissociation constant, and n is the Hill coefficient. All measurements were summarized in Excel (Microsoft) and imported into Prism (Graphpad Software Inc.) for statistical analysis and graph production.

### Immunoblot protocol

Immunoblots were performed as described previously, **(Vevea & Chapman, 2020)**, but were imaged on a ChemiDoc MP Imaging System (Biorad, 17001402). Primary and secondary antibodies are listed in the Key resource table with identifiers and dilutions used.

### Chemicals

Chemicals are listed in key resource table. All chemicals were resuspended and stored per manufacturer guidelines. Folimycin was ordered prior to use, stored dry at -20°C in original packaging, resuspended the week of the experiments and aliquots were frozen at -20°C, not to be used more than a week after resuspension.

### Statistics

Exact values from experiments and analysis, including number of data points (n) and number of trials, i.e. biological replicates, for each experiment are listed in the figure legends. GraphPad Prism 9.3.1 (GraphPad Software Inc.) was used for statistical analysis. When appropriate, data are displayed as Tukey box and whisker plots. Most data were not normally distributed, so nonparametric tests (i.e. Mann-Whitney test) were used throughout. Detailed statistical results for Figure 1d-f, Figure 2b-c, 2f-h, Figure 2 Figure supplement 1a, Figure 3b, 3f-i, Figure 3 Figure supplement 1a-c, Figure 4b-h, and 4j are included as source data files.

## Supplemental source and Video data legends

**Video 1. Asymmetric trafficking of mitochondria in axons**

Example image sequence of background-subtracted MTG+ and MTR+ axonal mitochondria trafficking through the microchannel of a 450 µm long microfluidic. The image sequence is an example from data collected for Figure 1c. Interval between frames are 5 sec, scale bar: 5 µm.

**Video 2. Anterograde mitochondria are more reduced and complement resident axonal mitochondria**

Example timeseries from the axon chamber of a microfluidic of mitochondria expressing roGFP, image sequence is the background-subtracted ratio of both emissions from roGFP. This is the full video from Figure 2d, where select images are shown. Fire LUT represents reduced (purple) or oxidized (white) signal, interval between frames is 5 sec. Scale bar: 5 µm.

**Video 3. Anterograde mitochondria are more reduced and complement resident axonal mitochondria**

Example image sequence of background-subtracted MTG+ and MTR+ axonal mitochondria trafficking in the axonal chamber, 2 hours post-MitoTracker label. The image sequence is an example from data collected for Figure 2e. Interval between frames are 5 sec, scale bar: 5 µm.

**Figure 1 – source data 1**

Descriptive statistics and statistical summary using two-tailed Mann-Whitney test analyzing mitochondrial length between anterograde and retrograde trafficking mitochondria.

**Figure 1 – source data 2**

Descriptive statistics and statistical summary using two-tailed Mann-Whitney test analyzing mitochondrial speed between anterograde and retrograde trafficking mitochondria.

**Figure 1 – source data 3**

Descriptive statistics and statistical summary using two-tailed Mann-Whitney test analyzing mitochondrial track length between anterograde and retrograde trafficking mitochondria.

**Figure 2 – source data 1**

Descriptive statistics and statistical summary using two-tailed Mann-Whitney test analyzing the normalized mitochondrial redox ratio between anterograde and retrograde trafficking mitochondria.

**Figure 2 – source data 2**

Descriptive statistics and statistical summary using two-tailed Mann-Whitney test analyzing mitochondrial length between anterograde and retrograde trafficking mitochondria.

**Figure 2 – source data 3**

Descriptive statistics summarizing the amount of new mitochondrial trafficking into the axon.

**Figure 2 – source data 4**

Descriptive statistics summarizing the fraction of new mitochondria (MTG) that fuse with resident axonal mitochondria, defined as the percentage of MTG that colocalized with MTR.

**Figure 2 – source data 5**

Descriptive statistics summarizing the fraction of resident axonal mitochondria (MTR) that fuse with newly trafficked mitochondria, defined as the percentage of MTR that colocalized with MTG.

**Figure 2 – Figure supplement 1 - source data 1**

Descriptive statistics and statistical summary using two-tailed Mann-Whitney test analyzing mitochondrial redox ratio between anterograde and retrograde trafficking mitochondria.

**Figure 3 – source data 1**

Statistical summary using Wilcoxon Signed Rank test analyzing the MFN2 protein level in MFN2 KD treated neurons.

**Figure 3 – source data 2**

Descriptive statistics and statistical summary using two-tailed Mann-Whitney test analyzing the total vesicle pool as measured by total vGlut1-pHluorin fluorescence between CTRL KD and MFN2 KD neurons.

**Figure 3 – source data 3**

Descriptive statistics and statistical summary using two-tailed Mann-Whitney test analyzing the endocytosis time constant (τ) between CTRL KD and MFN2 KD neurons.

**Figure 3 – source data 4**

Descriptive statistics and statistical summary using two-tailed Mann-Whitney test analyzing the total exocytosis between CTRL KD and MFN2 KD neurons.

**Figure 3 – source data 5**

Descriptive statistics and statistical summary using two-tailed Mann-Whitney test analyzing the total reserve pool of synaptic vesicles CTRL KD and MFN2 KD neurons.

**Figure 3 – Figure supplement 1 - source data 1**

Descriptive statistics and statistical summary using two-tailed Mann-Whitney test analyzing exocytosis between CTRL KD and MFN2 KD neurons.

**Figure 3 – Figure supplement 1 - source data 2**

Descriptive statistics and statistical summary using two-tailed Mann-Whitney test analyzing non-normalized exocytosis between CTRL KD and MFN2 KD neurons.

**Figure 3 – Figure supplement 1 - source data 3**

Descriptive statistics and statistical summary using two-tailed Mann-Whitney test analyzing non-normalized exocytosis between CTRL KD and MFN2 KD neurons.

**Figure 4 – source data 1**

Descriptive statistics and statistical summary using two-tailed Mann-Whitney test analyzing resting presynaptic Ca^2+^ between CTRL KD and MFN2 KD neurons.

**Figure 4 – source data 2**

Descriptive statistics and statistical summary using two-tailed Mann-Whitney test analyzing peak presynaptic Ca^2+^ after 1 AP, between CTRL KD and MFN2 KD neurons.

**Figure 4 – source data 3**

Descriptive statistics and statistical summary using two-tailed Mann-Whitney test analyzing the single-component time constant (τ) of Ca^2+^ decay after 1 AP, between CTRL KD and MFN2 KD neurons.

**Figure 4 – source data 4**

Descriptive statistics and values for the single-component best fit exponential decay after 1 AP, between CTRL KD and MFN2 KD neurons.

**Figure 4 – source data 5**

Descriptive statistics and statistical summary using two-tailed Mann-Whitney test analyzing the two-component time constants (τ) {FAST] and [SLOW] of Ca^2+^ decay after a 50 AP stimulus train at 50 Hz, between CTRL KD and MFN2 KD neurons.

**Figure 4 – source data 6**

Descriptive statistics and values for the two-component best fit exponential decay after 50 AP, between CTRL KD and MFN2 KD neurons.

**Figure 4 – source data 7**

Statistical summary using two-tailed Mann-Whitney test analyzing the single-component τ from 1 AP and the [FAST] τ from a 50 AP stimulus train, between CTRL KD and MFN2 KD neurons.

**Figure 4 – source data 8**

Descriptive statistics for proteins in the MFN2 KD treated neurons, normalized to CTRL KD.

## Notes

### Competing Interest Statement

The authors have declared no competing interest.

## References

Amiri, M., & Hollenbeck, P. J. (2008). Mitochondrial biogenesis in the axons of vertebrate peripheral neurons. Dev Neurobiol, 68(11), 1348–1361. doi:10.1002/dneu.20668

Area-Gomez, E., Guardia-Laguarta, C., Schon, E. A., & Przedborski, S. (2019). Mitochondria, OxPhos, and neurodegeneration: cells are not just running out of gas. J Clin Invest, 129(1), 34–45. doi:10.1172/JCI120848

Armbruster, M., Messa, M., Ferguson, S. M., De Camilli, P., & Ryan, T. A. (2013). Dynamin phosphorylation controls optimization of endocytosis for brief action potential bursts. Elife, 2, e00845. doi:10.7554/eLife.00845

Ashrafi, G., Schlehe, J. S., LaVoie, M. J., & Schwarz, T. L. (2014). Mitophagy of damaged mitochondria occurs locally in distal neuronal axons and requires PINK1 and Parkin. J Cell Biol, 206(5), 655–670. doi:10.1083/jcb.201401070

Atasoy, D., Ertunc, M., Moulder, K. L., Blackwell, J., Chung, C., Su, J., & Kavalali, E. T. (2008). Spontaneous and evoked glutamate release activates two populations of NMDA receptors with limited overlap. J Neurosci, 28(40), 10151–10166. doi:10.1523/JNEUROSCI.2432-08.2008

Balaji, J., Armbruster, M., & Ryan, T. A. (2008). Calcium control of endocytic capacity at a CNS synapse. J Neurosci, 28(26), 6742–6749. doi:10.1523/JNEUROSCI.1082-08.2008

Baloh, R. H., Schmidt, R. E., Pestronk, A., & Milbrandt, J. (2007). Altered axonal mitochondrial transport in the pathogenesis of Charcot-Marie-Tooth disease from mitofusin 2 mutations. J Neurosci, 27(2), 422–430. doi:10.1523/JNEUROSCI.4798-06.2007

Berthet, A., Margolis, E. B., Zhang, J., Hsieh, I., Zhang, J., Hnasko, T. S., … Nakamura, K. (2014). Loss of mitochondrial fission depletes axonal mitochondria in midbrain dopamine neurons. J Neurosci, 34(43), 14304–14317. doi:10.1523/JNEUROSCI.0930-14.2014

Billups, B., & Forsythe, I. D. (2002). Presynaptic mitochondrial calcium sequestration influences transmission at mammalian central synapses. J Neurosci, 22(14), 5840–5847. doi:20026597

Bomba-Warczak, E., Edassery, S. L., Hark, T. J., & Savas, J. N. (2021). Long-lived mitochondrial cristae proteins in mouse heart and brain. J Cell Biol, 220(9). doi:10.1083/jcb.202005193

Bomba-Warczak, E., Vevea, J. D., Brittain, J. M., Figueroa-Bernier, A., Tepp, W. H., Johnson, E. A., … Chapman, E. R. (2016). Interneuronal Transfer and Distal Action of Tetanus Toxin and Botulinum Neurotoxins A and D in Central Neurons. Cell Rep, 16(7), 1974–1987. doi:10.1016/j.celrep.2016.06.104

Bradberry, M. M., & Chapman, E. R. (2022). All-optical monitoring of excitation-secretion coupling demonstrates that SV2A functions downstream of evoked Ca(2+) entry. J Physiol, 600(3), 645–654. doi:10.1113/JP282601

Burrone, J., Li, Z., & Murthy, V. N. (2006). Studying vesicle cycling in presynaptic terminals using the genetically encoded probe synaptopHluorin. Nat Protoc, 1(6), 2970–2978. doi:10.1038/nprot.2006.449

Cai, Q., Zakaria, H. M., Simone, A., & Sheng, Z. H. (2012). Spatial parkin translocation and degradation of damaged mitochondria via mitophagy in live cortical neurons. Curr Biol, 22(6), 545–552. doi:10.1016/j.cub.2012.02.005

Chanaday, N. L., Cousin, M. A., Milosevic, I., Watanabe, S., & Morgan, J. R. (2019). The Synaptic Vesicle Cycle Revisited: New Insights into the Modes and Mechanisms. J Neurosci, 39(42), 8209–8216. doi:10.1523/JNEUROSCI.1158-19.2019

Chen, H., & Chan, D. C. (2009). Mitochondrial dynamics--fusion, fission, movement, and mitophagy--in neurodegenerative diseases. Hum Mol Genet, 18(R2), R169–176. doi:10.1093/hmg/ddp326

Chen, H., Chomyn, A., & Chan, D. C. (2005). Disruption of fusion results in mitochondrial heterogeneity and dysfunction. J Biol Chem, 280(28), 26185–26192. doi:10.1074/jbc.M503062200

Chen, H., Detmer, S. A., Ewald, A. J., Griffin, E. E., Fraser, S. E., & Chan, D. C. (2003). Mitofusins Mfn1 and Mfn2 coordinately regulate mitochondrial fusion and are essential for embryonic development. J Cell Biol, 160(2), 189–200. doi:10.1083/jcb.200211046

Cosson, P., Marchetti, A., Ravazzola, M., & Orci, L. (2012). Mitofusin-2 independent juxtaposition of endoplasmic reticulum and mitochondria: an ultrastructural study. PLoS One, 7(9), e46293. doi:10.1371/journal.pone.0046293

Craig, A. M., & Banker, G. (1994). Neuronal polarity. Annu Rev Neurosci, 17, 267–310. doi:10.1146/annurev.ne.17.030194.001411

Datta, S., & Jaiswal, M. (2021). Mitochondrial calcium at the synapse. Mitochondrion, 59, 135–153. doi:10.1016/j.mito.2021.04.006

de Brito, O. M., & Scorrano, L. (2008). Mitofusin 2 tethers endoplasmic reticulum to mitochondria. Nature, 456(7222), 605–610. doi:10.1038/nature07534

de Juan-Sanz, J., Holt, G. T., Schreiter, E. R., de Juan, F., Kim, D. S., & Ryan, T. A. (2017). Axonal Endoplasmic Reticulum Ca(2+) Content Controls Release Probability in CNS Nerve Terminals. Neuron, 93(4), 867–881 e866. doi:10.1016/j.neuron.2017.01.010

Delvendahl, I., Vyleta, N. P., von Gersdorff, H., & Hallermann, S. (2016). Fast, Temperature-Sensitive and Clathrin-Independent Endocytosis at Central Synapses. Neuron, 90(3), 492–498. doi:10.1016/j.neuron.2016.03.013

Deo, C., Sheu, S. H., Seo, J., Clapham, D. E., & Lavis, L. D. (2019). Isomeric Tuning Yields Bright and Targetable Red Ca(2+) Indicators. J Am Chem Soc, 141(35), 13734–13738. doi:10.1021/jacs.9b06092

Dooley, C. T., Dore, T. M., Hanson, G. T., Jackson, W. C., Remington, S. J., & Tsien, R. Y. (2004). Imaging dynamic redox changes in mammalian cells with green fluorescent protein indicators. J Biol Chem, 279(21), 22284–22293. doi:10.1074/jbc.M312847200

Ebrahimi-Fakhari, D., Saffari, A., Wahlster, L., DiNardo, A., Turner, D., Lewis, T. L., Jr., … Sahin, M. (2016). Impaired Mitochondrial Dynamics And Mitophagy In Neuronal Models Of Tuberous Sclerosis Complex. Cell Rep, 17(8), 2162. doi:10.1016/j.celrep.2016.10.051

Eisner, V., Picard, M., & Hajnoczky, G. (2018). Mitochondrial dynamics in adaptive and maladaptive cellular stress responses. Nat Cell Biol, 20(7), 755–765. doi:10.1038/s41556-018-0133-0

Eura, Y., Ishihara, N., Yokota, S., & Mihara, K. (2003). Two mitofusin proteins, mammalian homologues of FZO, with distinct functions are both required for mitochondrial fusion. J Biochem, 134(3), 333–344. doi:10.1093/jb/mvg150

Evans, C. S., & Holzbaur, E. L. (2020). Degradation of engulfed mitochondria is rate-limiting in Optineurin-mediated mitophagy in neurons. Elife, 9. doi:10.7554/eLife.50260

Faitg, J., Lacefield, C., Davey, T., White, K., Laws, R., Kosmidis, S., … Picard, M. (2021). 3D neuronal mitochondrial morphology in axons, dendrites, and somata of the aging mouse hippocampus. Cell Rep, 36(6), 109509. doi:10.1016/j.celrep.2021.109509

Fang, D., Yan, S., Yu, Q., Chen, D., & Yan, S. S. (2016). Mfn2 is Required for Mitochondrial Development and Synapse Formation in Human Induced Pluripotent Stem Cells/hiPSC Derived Cortical Neurons. Sci Rep, 6, 31462. doi:10.1038/srep31462

Filadi, R., Greotti, E., Turacchio, G., Luini, A., Pozzan, T., & Pizzo, P. (2015). Mitofusin 2 ablation increases endoplasmic reticulum-mitochondria coupling. Proc Natl Acad Sci U S A, 112(17), E2174–2181. doi:10.1073/pnas.1504880112

Hanson, G. T., Aggeler, R., Oglesbee, D., Cannon, M., Capaldi, R. A., Tsien, R. Y., & Remington, S. J. (2004). Investigating mitochondrial redox potential with redox-sensitive green fluorescent protein indicators. J Biol Chem, 279(13), 13044–13053. doi:10.1074/jbc.M312846200

Higuchi, R., Vevea, J. D., Swayne, T. C., Chojnowski, R., Hill, V., Boldogh, I. R., & Pon, L. A. (2013). Actin dynamics affect mitochondrial quality control and aging in budding yeast. Curr Biol, 23(23), 2417–2422. doi:10.1016/j.cub.2013.10.022

Ishihara, N., Nomura, M., Jofuku, A., Kato, H., Suzuki, S. O., Masuda, K., … Mihara, K. (2009). Mitochondrial fission factor Drp1 is essential for embryonic development and synapse formation in mice. Nat Cell Biol, 11(8), 958–966. doi:10.1038/ncb1907

Jordan, M., & Wurm, F. (2004). Transfection of adherent and suspended cells by calcium phosphate. Methods, 33(2), 136–143. doi:10.1016/j.ymeth.2003.11.011

Katajisto, P., Dohla, J., Chaffer, C. L., Pentinmikko, N., Marjanovic, N., Iqbal, S., … Sabatini, D. M. (2015). Stem cells. Asymmetric apportioning of aged mitochondria between daughter cells is required for stemness. Science, 348(6232), 340–343. doi:10.1126/science.1260384

Koester, H. J., & Sakmann, B. (2000). Calcium dynamics associated with action potentials in single nerve terminals of pyramidal cells in layer 2/3 of the young rat neocortex. J Physiol, 529 Pt 3, 625–646. doi:10.1111/j.1469-7793.2000.00625.x

Kowaltowski, A. J., Menezes-Filho, S. L., Assali, E. A., Goncalves, I. G., Cabral-Costa, J. V., Abreu, P., … Shirihai, O. S. (2019). Mitochondrial morphology regulates organellar Ca(2+) uptake and changes cellular Ca(2+) homeostasis. FASEB J, 33(12), 13176–13188. doi:10.1096/fj.201901136R

Kugler, S., Kilic, E., & Bahr, M. (2003). Human synapsin 1 gene promoter confers highly neuron-specific long-term transgene expression from an adenoviral vector in the adult rat brain depending on the transduced area. Gene Ther, 10(4), 337–347. doi:10.1038/sj.gt.3301905

Kuzniewska, B., Cysewski, D., Wasilewski, M., Sakowska, P., Milek, J., Kulinski, T. M., … Dziembowska, M. (2020). Mitochondrial protein biogenesis in the synapse is supported by local translation. EMBO Rep, 21(8), e48882. doi:10.15252/embr.201948882

Leitz, J., & Kavalali, E. T. (2011). Ca(2)(+) influx slows single synaptic vesicle endocytosis. J Neurosci, 31(45), 16318–16326. doi:10.1523/JNEUROSCI.3358-11.2011

Leitz, J., & Kavalali, E. T. (2016). Ca2+ Dependence of Synaptic Vesicle Endocytosis. Neuroscientist, 22(5), 464–476. doi:10.1177/1073858415588265

Lewis, T. L., Jr., Kwon, S. K., Lee, A., Shaw, R., & Polleux, F. (2018). MFF-dependent mitochondrial fission regulates presynaptic release and axon branching by limiting axonal mitochondria size. Nat Commun, 9(1), 5008. doi:10.1038/s41467-018-07416-2

Li, Z., Burrone, J., Tyler, W. J., Hartman, K. N., Albeanu, D. F., & Murthy, V. N. (2005). Synaptic vesicle recycling studied in transgenic mice expressing synaptopHluorin. Proc Natl Acad Sci U S A, 102(17), 6131–6136. doi:10.1073/pnas.0501145102

Liao, P. C., Tandarich, L. C., & Hollenbeck, P. J. (2017). ROS regulation of axonal mitochondrial transport is mediated by Ca2+ and JNK in Drosophila. PLoS One, 12(5), e0178105. doi:10.1371/journal.pone.0178105

Lin, M. Y., Cheng, X. T., Tammineni, P., Xie, Y., Zhou, B., Cai, Q., & Sheng, Z. H. (2017). Releasing Syntaphilin Removes Stressed Mitochondria from Axons Independent of Mitophagy under Pathophysiological Conditions. Neuron, 94(3), 595–610 e596. doi:10.1016/j.neuron.2017.04.004

Lin, T. H., Bis-Brewer, D. M., Sheehan, A. E., Townsend, L. N., Maddison, D. C., Zuchner, S., … Freeman, M. R. (2021). TSG101 negatively regulates mitochondrial biogenesis in axons. Proc Natl Acad Sci U S A, 118(20). doi:10.1073/pnas.2018770118

Lois, C., Hong, E. J., Pease, S., Brown, E. J., & Baltimore, D. (2002). Germline transmission and tissue-specific expression of transgenes delivered by lentiviral vectors. Science, 295(5556), 868–872. doi:10.1126/science.1067081

Mandal, A., Wong, H. C., Pinter, K., Mosqueda, N., Beirl, A., Lomash, R. M., … Drerup, C. M. (2021). Retrograde Mitochondrial Transport Is Essential for Organelle Distribution and Health in Zebrafish Neurons. J Neurosci, 41(7), 1371–1392. doi:10.1523/JNEUROSCI.1316-20.2020

Maravall, M., Mainen, Z. F., Sabatini, B. L., & Svoboda, K. (2000). Estimating intracellular calcium concentrations and buffering without wavelength ratioing. Biophys J, 78(5), 2655–2667. doi:10.1016/S0006-3495(00)76809-3

Marland, J. R., Hasel, P., Bonnycastle, K., & Cousin, M. A. (2016). Mitochondrial Calcium Uptake Modulates Synaptic Vesicle Endocytosis in Central Nerve Terminals. J Biol Chem, 291(5), 2080–2086. doi:10.1074/jbc.M115.686956

Matsuda, T., & Cepko, C. L. (2004). Electroporation and RNA interference in the rodent retina in vivo and in vitro. Proc Natl Acad Sci U S A, 101(1), 16–22. doi:10.1073/pnas.2235688100

Mattie, S., Riemer, J., Wideman, J. G., & McBride, H. M. (2018). A new mitofusin topology places the redox-regulated C terminus in the mitochondrial intermembrane space. J Cell Biol, 217(2), 507–515. doi:10.1083/jcb.201611194

McFaline-Figueroa, J. R., Vevea, J., Swayne, T. C., Zhou, C., Liu, C., Leung, G., … Pon, L. A. (2011). Mitochondrial quality control during inheritance is associated with lifespan and mother-daughter age asymmetry in budding yeast. Aging Cell, 10(5), 885–895. doi:10.1111/j.1474-9726.2011.00731.x

Miesenbock, G., De Angelis, D. A., & Rothman, J. E. (1998). Visualizing secretion and synaptic transmission with pH-sensitive green fluorescent proteins. Nature, 394(6689), 192–195. doi:10.1038/28190

Miller, K. E., & Sheetz, M. P. (2004). Axonal mitochondrial transport and potential are correlated. J Cell Sci, 117(Pt 13), 2791–2804. doi:10.1242/jcs.01130

Misko, A., Jiang, S., Wegorzewska, I., Milbrandt, J., & Baloh, R. H. (2010). Mitofusin 2 is necessary for transport of axonal mitochondria and interacts with the Miro/Milton complex. J Neurosci, 30(12), 4232–4240. doi:10.1523/JNEUROSCI.6248-09.2010

Misko, A. L., Sasaki, Y., Tuck, E., Milbrandt, J., & Baloh, R. H. (2012). Mitofusin2 mutations disrupt axonal mitochondrial positioning and promote axon degeneration. J Neurosci, 32(12), 4145–4155. doi:10.1523/JNEUROSCI.6338-11.2012

Nakada, K., Inoue, K., Ono, T., Isobe, K., Ogura, A., Goto, Y. I., … Hayashi, J. I. (2001). Inter-mitochondrial complementation: Mitochondria-specific system preventing mice from expression of disease phenotypes by mutant mtDNA. Nat Med, 7(8), 934–940. doi:10.1038/90976

Naon, D., Zaninello, M., Giacomello, M., Varanita, T., Grespi, F., Lakshminaranayan, S., … Scorrano, L. (2016). Critical reappraisal confirms that Mitofusin 2 is an endoplasmic reticulum-mitochondria tether. Proc Natl Acad Sci U S A, 113(40), 11249–11254. doi:10.1073/pnas.1606786113

Neher, E., & Zucker, R. S. (1993). Multiple calcium-dependent processes related to secretion in bovine chromaffin cells. Neuron, 10(1), 21–30. doi:10.1016/0896-6273(93)90238-m

Neves, G., Gomis, A., & Lagnado, L. (2001). Calcium influx selects the fast mode of endocytosis in the synaptic terminal of retinal bipolar cells. Proc Natl Acad Sci U S A, 98(26), 15282–15287. doi:10.1073/pnas.261311698

Oettinghaus, B., Schulz, J. M., Restelli, L. M., Licci, M., Savoia, C., Schmidt, A., … Frank, S. (2016). Synaptic dysfunction, memory deficits and hippocampal atrophy due to ablation of mitochondrial fission in adult forebrain neurons. Cell Death Differ, 23(1), 18–28. doi:10.1038/cdd.2015.39

Overly, C. C., Rieff, H. I., & Hollenbeck, P. J. (1996). Organelle motility and metabolism in axons vs dendrites of cultured hippocampal neurons. J Cell Sci, 109 (Pt 5), 971–980. doi:10.1242/jcs.109.5.971

Patterson, G. H., & Lippincott-Schwartz, J. (2002). A photoactivatable GFP for selective photolabeling of proteins and cells. Science, 297(5588), 1873–1877. doi:10.1126/science.1074952

Pickles, S., Vigie, P., & Youle, R. J. (2018). Mitophagy and Quality Control Mechanisms in Mitochondrial Maintenance. Curr Biol, 28(4), R170–R185. doi:10.1016/j.cub.2018.01.004

Regehr, W. G., & Atluri, P. P. (1995). Calcium transients in cerebellar granule cell presynaptic terminals. Biophys J, 68(5), 2156–2170. doi:10.1016/S0006-3495(95)80398-X

Rojo, M., Legros, F., Chateau, D., & Lombes, A. (2002). Membrane topology and mitochondrial targeting of mitofusins, ubiquitous mammalian homologs of the transmembrane GTPase Fzo. J Cell Sci, 115(Pt 8), 1663–1674. doi:10.1242/jcs.115.8.1663

Sankaranarayanan, S., De Angelis, D., Rothman, J. E., & Ryan, T. A. (2000). The use of pHluorins for optical measurements of presynaptic activity. Biophys J, 79(4), 2199–2208. doi:10.1016/S0006-3495(00)76468-X

Sankaranarayanan, S., & Ryan, T. A. (2000). Real-time measurements of vesicle-SNARE recycling in synapses of the central nervous system. Nat Cell Biol, 2(4), 197–204. doi:10.1038/35008615

Sankaranarayanan, S., & Ryan, T. A. (2001). Calcium accelerates endocytosis of vSNAREs at hippocampal synapses. Nat Neurosci, 4(2), 129–136. doi:10.1038/83949

Saxton, W. M., & Hollenbeck, P. J. (2012). The axonal transport of mitochondria. J Cell Sci, 125(Pt 9), 2095–2104. doi:10.1242/jcs.053850

Schon, E. A., & Przedborski, S. (2011). Mitochondria: the next (neurode)generation. Neuron, 70(6), 1033–1053. doi:10.1016/j.neuron.2011.06.003

Shutt, T., Geoffrion, M., Milne, R., & McBride, H. M. (2012). The intracellular redox state is a core determinant of mitochondrial fusion. EMBO Rep, 13(10), 909–915. doi:10.1038/embor.2012.128

Spinelli, J. B., & Haigis, M. C. (2018). The multifaceted contributions of mitochondria to cellular metabolism. Nat Cell Biol, 20(7), 745–754. doi:10.1038/s41556-018-0124-1

Sun, J. Y., Wu, X. S., & Wu, L. G. (2002). Single and multiple vesicle fusion induce different rates of endocytosis at a central synapse. Nature, 417(6888), 555–559. doi:10.1038/417555a

Suzuki, R., Hotta, K., & Oka, K. (2018). Transitional correlation between inner-membrane potential and ATP levels of neuronal mitochondria. Sci Rep, 8(1), 2993. doi:10.1038/s41598-018-21109-2

Tank, D. W., Regehr, W. G., & Delaney, K. R. (1995). A quantitative analysis of presynaptic calcium dynamics that contribute to short-term enhancement. J Neurosci, 15(12), 7940–7952. Retrieved from https://www.ncbi.nlm.nih.gov/pubmed/8613732

Trevisan, T., Pendin, D., Montagna, A., Bova, S., Ghelli, A. M., & Daga, A. (2018). Manipulation of Mitochondria Dynamics Reveals Separate Roles for Form and Function in Mitochondria Distribution. Cell Rep, 23(6), 1742–1753. doi:10.1016/j.celrep.2018.04.017

Twig, G., Elorza, A., Molina, A. J., Mohamed, H., Wikstrom, J. D., Walzer, G., … Shirihai, O. S. (2008). Fission and selective fusion govern mitochondrial segregation and elimination by autophagy. EMBO J, 27(2), 433–446. doi:10.1038/sj.emboj.7601963

Verburg, J., & Hollenbeck, P. J. (2008). Mitochondrial membrane potential in axons increases with local nerve growth factor or semaphorin signaling. J Neurosci, 28(33), 8306–8315. doi:10.1523/JNEUROSCI.2614-08.2008

Verstreken, P., Ly, C. V., Venken, K. J., Koh, T. W., Zhou, Y., & Bellen, H. J. (2005). Synaptic mitochondria are critical for mobilization of reserve pool vesicles at Drosophila neuromuscular junctions. Neuron, 47(3), 365–378. doi:10.1016/j.neuron.2005.06.018

Vevea, J. D., Alessi Wolken, D. M., Swayne, T. C., White, A. B., & Pon, L. A. (2013). Ratiometric biosensors that measure mitochondrial redox state and ATP in living yeast cells. J Vis Exp(77), 50633. doi:10.3791/50633

Vevea, J. D., & Chapman, E. R. (2020). Acute disruption of the synaptic vesicle membrane protein synaptotagmin 1 using knockoff in mouse hippocampal neurons. Elife, 9. doi:10.7554/eLife.56469

Voglmaier, S. M., Kam, K., Yang, H., Fortin, D. L., Hua, Z., Nicoll, R. A., & Edwards, R. H. (2006). Distinct endocytic pathways control the rate and extent of synaptic vesicle protein recycling. Neuron, 51(1), 71–84. doi:10.1016/j.neuron.2006.05.027

von Gersdorff, H., & Matthews, G. (1994). Inhibition of endocytosis by elevated internal calcium in a synaptic terminal. Nature, 370(6491), 652–655. doi:10.1038/370652a0

Wang, X., Winter, D., Ashrafi, G., Schlehe, J., Wong, Y. L., Selkoe, D., … Schwarz, T. L. (2011). PINK1 and Parkin target Miro for phosphorylation and degradation to arrest mitochondrial motility. Cell, 147(4), 893–906. doi:10.1016/j.cell.2011.10.018

Werth, J. L., & Thayer, S. A. (1994). Mitochondria buffer physiological calcium loads in cultured rat dorsal root ganglion neurons. J Neurosci, 14(1), 348–356. Retrieved from https://www.ncbi.nlm.nih.gov/pubmed/8283242

Wu, W., Xu, J., Wu, X. S., & Wu, L. G. (2005). Activity-dependent acceleration of endocytosis at a central synapse. J Neurosci, 25(50), 11676–11683. doi:10.1523/JNEUROSCI.2972-05.2005

Zhang, W., & Linden, D. J. (2012). Calcium influx measured at single presynaptic boutons of cerebellar granule cell ascending axons and parallel fibers. Cerebellum, 11(1), 121–131. doi:10.1007/s12311-009-0151-3

Zheng, Y., Zhang, X., Wu, X., Jiang, L., Ahsan, A., Ma, S., … Chen, Z. (2019). Somatic autophagy of axonal mitochondria in ischemic neurons. J Cell Biol, 218(6), 1891–1907. doi:10.1083/jcb.201804101

